# The FORGENIUS genomic resources: new genotyping tools and genomic data for 23 forest tree species and their Genetic Conservation Units

**DOI:** 10.1101/2025.08.08.669074

**Authors:** Sara Pinosio, Francesca Bagnoli, Camilla Avanzi, Maria B. Castellani, Arcangela Frascella, Susan L. McEvoy, Sanna Olsson, Ilaria Spanu, Elia Vajana, the FORGENIUS Consortium, Santiago C. González-Martínez, Tanja Pyhäjärvi, Ivan Scotti, Giovanni G. Vendramin, Andrea Piotti

## Abstract

Genetic diversity is a critical but often overlooked component of biological diversity. The European H2020 FORGENIUS project is precisely aimed at increasing the quality and quantity of genetic data to start monitoring the European network of forest Genetic Conservation Units (GCUs). A first step in this direction was developing standardized genomic resources for 23 forest tree species, spanning from rare and scattered (e.g., *Abies nebrodensis* and *Torminalis glaberrima*) to widespread, economically relevant ones (e.g., *Fagus sylvatica, Picea abies* and *Pinus sylvestris*).

Here, we describe the development and application of targeted genotyping tools, primarily based on Single Primer Enrichment Technology (SPET), along with existing SNP arrays for the selected species. The SPET panels developed in FORGENIUS were designed to capture □10,000 loci per species, balancing species-specific and randomly distributed regions to ensure broad genome coverage and minimize ascertainment bias.

Across 7,192 genotyped trees, we identified over 1.8 million single nucleotide polymorphisms (SNPs) covering approximately 50 Mb of DNA sequence. SPET panels demonstrated high genotyping efficiency and cross-species transferability, especially within genera such as *Quercus* and *Abies*. They represent a cost-effective, flexible, and scalable solution for population-level genetic assessments across diverse taxa, enabling standardized, genome-wide characterization of the GCU network. These resources not only promote the establishment of genetic monitoring, support genetically informed conservation strategies and improve our understanding of adaptive responses in European forests, but also enhance species delimitation and hybrid detection, and enable the characterization of phylogenetically related but previously underexplored species.

## Introduction

Despite being a key dimension of biodiversity, genetic diversity has been long overlooked in international conservation programs (Laikre et al., 2020) until very recent initiatives (Hoban et al., 2025). Up to 16% of global genetic diversity is expected to go extinct by the end of the 21^st^ century due to habitat loss (Exposito-Alonso et al., 2022) and proper genomic tools are crucial to track its future changes (Pearman et al., 2024; Shaw et al., 2025). Genetic monitoring must rely on population surveys or inventories that can be repeated through time. To this aim, an asset for European forests is the continent-wide network of *in situ* Genetic Conservation Units (GCUs) coordinated by the European Forest Genetic Resources Programme (EUFORGEN) since 1994 (Lefèvre et al., 2013) and accessible through the EUFGIS information system (http://portal.eufgis.org). The GCU network is designed to represent forest tree stands that are adapted to unique sets of environmental conditions and are thus expected to have distinct genetic, phenotypic, and/or ecological characteristics.

Decisions on the inclusion of GCUs into the EUFORGEN collection have mostly been based on non-standardized local observations, rather than on quantitative scientific assessments (de Vries et al., 2015). For instance, genetic information is lacking in the EUFGIS information system, as well as information about structural, physiological and climatic characteristics of the forest stands in the GCUs. In addition, the proxy used to identify differently adapted GCUs (i.e. climatic zonation) omits many other potential drivers of natural selection and is far from optimal. The urgent need for a deep, functional characterization of GCUs requires information about adaptive variation at the molecular and phenotypic levels (EUFORGEN, 2021), and its integration with environmental data. Although a few, recently published studies include both a range- and genome-wide characterization of Europe’s most iconic tree species (e.g Bruxaux et al., 2024; Milesi et al., 2024; Theraroz et al., 2024; Zhou et al., 2024), much of the genetic information on the few genetically characterized European forest tree species is still based on a limited number of neutral molecular markers (Aravanopoulos, et al., 2015) that does not adequately describe their genetic diversity, in general, and, in particular, their adaptive potential. Since the advent of high-throughput sequencing techniques, however, it is possible to apply the same genotyping standard—sequencing itself or sequence-based genotyping—to all species, making genomic information homogeneous in kind (if not in completeness). Indeed, one of the overarching goals of the European H2020 project FORGENIUS (http://www.forgenius.eu) is to address the lack of knowledge on the European GCU network and provide a common baseline for monitoring the levels of neutral and adaptive genetic variation in European forests through time.

Several techniques based on next-generation sequencing (such as RAD-Seq, exome capture, and Pool-Seq) or DNA arrays can be adopted for rapid and cost-effective genomic characterization of natural resources. The targeted genotyping system based on Single Primer Enrichment Technology (SPET, Scaglione et al., 2019) is one of the most promising sequencing-based approaches and is increasingly used to characterize the genetic diversity of forest tree species (Budde et al., 2024; Olsson et al., 2023). SPET uses a single primer to selectively enrich specific regions of the genome, enabling targeted genotyping with reduced sequencing costs. This approach efficiently targets single nucleotide polymorphisms (SNPs) of interest while also yielding hundreds of thousands of untargeted variable sites (referred to as target and *de novo* SNPs, respectively). The sequencing read length can be adjusted based on the need to detect unknown (*de novo*) SNPs or to focus on cost-effective target genotyping. Additionally, the possibility to target randomly distributed regions of the genome ensures the generation of ascertainment bias-free markers. SPET panels can be easily transferred to other populations or even species and applied across different laboratories, facilitating the integration of data from multiple sources. The high reproducibility of SPET data makes it an excellent tool for population genomic analysis of natural populations and genetic monitoring.

Here, we describe the genomic resources developed and data produced for 23 forest tree species within the European H2020 FORGENIUS project. Five species were genotyped using existing, well-established tools based on array technology, while for all the other species we developed new SPET panels. These panels were designed to target approximately 10,000 genomic loci, uniformly distributed across the genome, combining both randomly selected regions and species-specific regions of interest. Using SPET panels and SNP array chips, we identified a total of 1,852,293 SNPs across 7,192 trees, covering about 50 Mb of genomic sequence. We describe the development of these genomic resources, from the design of SPET panels to data production, highlighting the different strategies adopted to obtain equally high-quality markers despite species-to-species variation in properties, such as presence/absence of a reference genome or proportion of repeated elements in the genome (which varies considerably between angiosperms and gymnosperms). We present overall results of genetic diversity and structure at the species level, emphasizing the importance of including a fraction of randomly selected loci in the set of target genomic regions, to ensure that estimates are unaffected by ascertainment bias. Finally, we discuss the relevance of the FORGENIUS genomic resources for the genomic characterization of the European network of GCUs, as well as their broader applications, ranging from genetic monitoring and the assessment of genomic vulnerability to the study of within-population dynamics (gene flow, hybridization, sexual vs vegetative reproduction, etc.) and the reconstruction of past evolutionary history.

## Materials and Methods

### Genomic characterization of forest Genetic Conservation Units (GCUs)

Twenty-three forest tree species were selected within FORGENIUS to provide a detailed and in-depth characterisation of a representative subset of the GCU network, as they globally occur in ⍰80% of GCUs. The selected species range from some of the most widespread ecologically and economically relevant in Europe (e.g., *Fagus sylvatica, Pinus sylvestris, Picea abies, Quercus petraea*) to rare species, such as *Abies nebrodensis*, or scattered species, such as *Malus sylvestris* and *Torminalis glaberrima* (formerly *Sorbus torminalis*). They cover a suite of environmental requirements, are present in most European eco-regions, and correspond to multiple societal demands (Figure 1).

**Figure 1.**
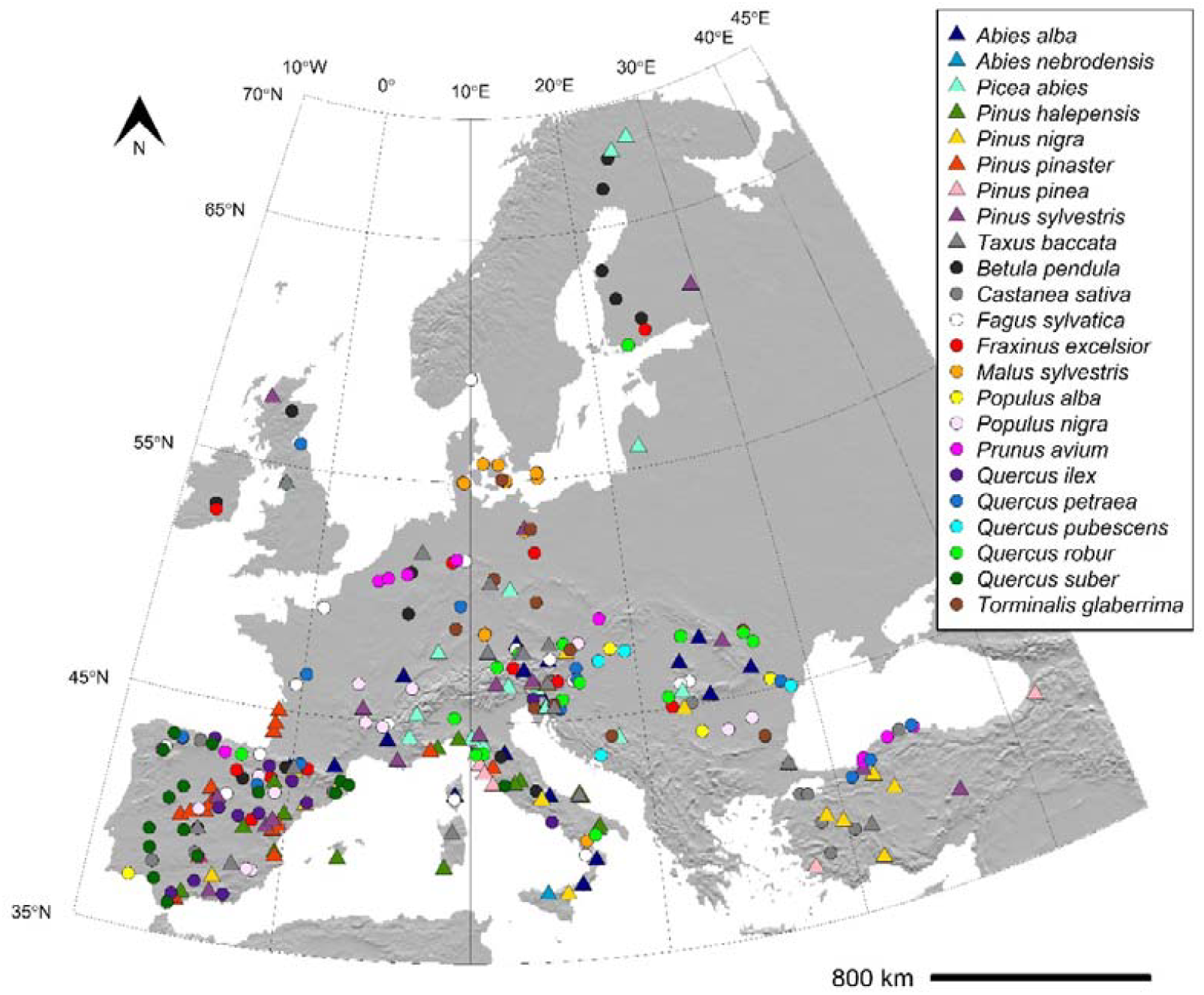
Distribution of the 300 GCUs of 23 forest tree species sampled for genomic characterization (details in Table S1). Different symbols refer to conifers (triangles) and broad-leaved species (circles).

Within each species, GCUs were selected to cover all environmental zones in which they occur, as well as to include ecologically and geographically marginal populations. The number of GCUs characterized per species spans from one (the only existing population of *A. nebrodensis*) to 18 (*Abies alba*) and is approximately proportional to the total number of GCUs for the focal species, totalling 300 GCUs for which genomic data were generated (Figure 1, Table S1 in the Supplementary Materials).

### Single Primer Enrichment Technology (SPET) panels design

Species-specific SPET probes were custom designed for fourteen species on the latest version of their reference genome, if available, or on the transcriptome sequence. Details on the reference sequences used for probe design are provided in Table 1.

**Table 1.** List of reference genomes or transcriptomes used for probe design and for data analysis.

| Species | Reference for probe design | Reference for data analysis |
| --- | --- | --- |
| *Abies alba* | *Abies alba* Abal.1_1 genome (Mosca et al., 2019) | *Abies alba* Abal.1_1 genome (Mosca et al., 2019) |
| *Betula pendula* | *Betula pendula* Bpev01 genome (Salojärvi et al., 2017) | *Betula pendula* Bpev01 genome (Salojärvi et al., 2017) |
| *Castanea sativa* | *Castanea sativa* transcriptome (Acquadro et al., 2019) | *Castanea mollissima* Mahogany genome (http://phytozome.jgi.doe.gov/info/CmollissimaMahoganyHAP1_v1_1) |
| *Fagus sylvatica* | *Fagus sylvatica* v2 genome (Mishra et al., 2022) | *Fagus sylvatica* v2 genome (Mishra et al., 2022) |
| *Fraxinus excelsior* | *Fraxinus excelsior* FRAX-001 genome (PRJNA713541) | *Fraxinus excelsior* FRAX-001 genome (PRJNA713541) |
| *Malus sylvestris* | *Malus sylvestris* haploid_v2 genome (PRJNA599188) | *Malus sylvestris* haploid_v2 genome (PRJNA599188) |
| *Picea abies* | *Picea abies* 1.0 genome (Nystedt et al., 2013) | *Picea abies* 1.0 genome (Nystedt et al., 2013) |
| *Pinus halepensis* | *Pinus halepensis* transcriptome (Pinosio et al., 2014) | *Pinus tabulaeformis* genome v1.0 (Kesälahti et al., 2025) |
| *Pinus nigra* | *Pinus nigra* transcriptome (Olsson et al., 2020) | *Pinus tabulaeformis* genome v1.0 (Kesälahti et al., 2025) |
| *Prunus avium* | *Prunus avium* Tieton_v2.0 genome (Wang et al., 2020) | *Prunus avium* Tieton_v2.0 genome (Wang et al., 2020) |
| *Quercus ilex* | *Quercus ilex* MR-2023 genome (Rey et al., 2023) | *Quercus ilex* MR-2023 genome (Rey et al., 2023) |
| *Quercus robur* | *Quercus robur* PM1N genome (Plomion et al., 2018) | *Quercus robur* PM1N genome (Plomion et al., 2018) |
| *Taxus baccata* | *Taxus baccata* transcriptome (Olsson et al., 2018) | *Taxus chinensis* genome (Xiong et al., 2021) |
| *Torminalis glaberrima* | *Torminalis glaberrima* transcriptome (Pinosio et al., 2025) | *Sorbus pohuashanensis* genome (Zhao et al., 2022) |

For all species genotyped by SPET, at least half of the probes were designed to capture random sites of the genome/transcriptome to obtain a set of SNPs without ascertainment bias. In the species for which molecular markers were already available, probes targeting known SNPs were also included. For each species, excluding *Quercus robur* and *F. sylvatica*, the corresponding reference genome or transcriptome, along with a BED file containing the selected target SNPs and random sites, were submitted to Tecan Genomics (Männedorf, Zurich, Switzerland) for probe design. For *Q. robur* and *F. sylvatica*, probes targeting both random sites and SNPs were selected from two pre-existing 90K SPET panels (Budde et al., 2024; Tost et al., 2025). A total of four distinct SPET probe panels were developed: 1) a species-specific panel including 10,000 probes for *A. alba* (FORGENIUS-Aalb10K), 2) a species-specific panel including 10,000 probes for *Pinus halepensis* (FORGENIUS-Phal10K), 3) a multi-species panel comprising 60,000 probes (FORGENIUS-MultiSp60K) with 10,000 probes designed for each of the six included species (i.e., *Betula pendula, Fraxinus excelsior, M. sylvestris, Pinus nigra, Prunus avium*, and *Taxus baccata*), and 4) a multi-species panel comprising 53,500 probes (FORGENIUS-MultiSp50K), including 10,000 probes for *Castanea sativa, F. sylvatica, P. abies, Q. robur*, and *T. glaberrima*, along with 3,500 probes for *Quercus ilex*. These FORGENIUS SPET panels are available at IGATech (https://igatechnology.com/sequencing-services/targeted-genotyping/). Supplementary Table 2 provides detailed information on each panel, including probe sequences, target site positions, and site type (i.e., target SNP or random site).

**Table 2.** Summary of sequencing metrics for each species included in the study. The table reports the average number of sequenced read pairs per sample, the percentage of uniquely aligned reads, the average insert size, the number of working probes, the mean depth at target regions, and the total megabases (Mb) covered across the target space.

| Species | Sequenced read pairs | % Aligned reads | Insert size | Nr. of working probes | Depth at target regions | Mb Covered |
| --- | --- | --- | --- | --- | --- | --- |
| *Abies alba* | 2,790,277 | 87.6 | 261 | 9,627 | 39.3 | 1.94 |
| *Abies nebrodensis* | 2,457,400 | 80.7 | 255 | 9,291 | 18.3 | 1.23 |
| *Betula pendula* | 1,937,435 | 76.0 | 308 | 7,735 | 61.3 | 2.35 |
| *Castanea sativa* | 1,349,141 | 88.0 | 254 | 14,004 | 55.6 | 3.56 |
| *Fagus sylvatica* | 1,193,929 | 83.1 | 286 | 11,446 | 53.5 | 3.02 |
| *Fraxinus excelsior* | 1,887,624 | 91.2 | 331 | 9,624 | 85.1 | 3.78 |
| *Malus sylvestris* | 1,706,058 | 86.0 | 297 | 9,139 | 62.1 | 2.89 |
| *Picea abies* | 2,733,083 | 32.5 | 256 | 9,105 | 21.2 | 1.12 |
| *Pinus halepensis* | 1,795,163 | 80.9 | 341 | 7,594 | 22.5 | 1.21 |
| *Pinus nigra* | 3,451,786 | 65.0 | 272 | 9,380 | 35.0 | 1.53 |
| *Prunus avium* | 1,283,333 | 89.7 | 390 | 9,456 | 41.0 | 3.52 |
| *Quercus ilex* | 1,290,987 | 87.7 | 225 | 10,631 | 58.3 | 2.43 |
| *Quercus petraea* | 2,084,643 | 86.3 | 277 | 16,511 | 60.6 | 4.39 |
| *Quercus pubescens* | 1,627,385 | 78.9 | 291 | 16,207 | 36.7 | 3.59 |
| *Quercus robur* | 2,803,723 | 86.3 | 308 | 16,657 | 51.0 | 4.38 |
| *Quercus suber* | 1,264,732 | 86.4 | 293 | 9,861 | 50.3 | 2.52 |
| *Taxus baccata* | 2,281,221 | 75.0 | 258 | 9,485 | 59.7 | 2.61 |
| *Torminalis glaberrima* | 2,617,148 | 91.9 | 199 | 9,717 | 149.9 | 2.74 |

### Selection of random target sites for probe design on reference genomes and transcriptomes

Whenever a reference genome was available, probes targeting random sites of the genome were chosen on non-repetitive regions using a k-mer approach. To this aim, Tallymer tool (Kurtz et al., 2008) included in the GenomeTools suite version 1.4.1 (Gremme et al., 2013), was run to estimate the mappability along the genome by calculating the number of occurrences of each genomic 20-mer. In detail, the Tallymer subroutine *suffixerator* was run with options -dna -pl -tis -suf -lcp -v -parts 16, while the subroutine *mkindex* was run with options -mersize 20 -minocc 1 -pl -counts. Thus, random positions were selected in regions at least 200 bp long and characterized by an average 20-mer value lower than 2.

When a reference genome was unavailable and a transcriptome was used instead, the genome of a related species was used to identify regions with high mappability that were also contiguous between the transcriptome and genome. This approach ensured that probes were not designed across exon-intron junctions or within repetitive sequences. To this aim, the *wgsim* utility included in the samtools v1.7 package (Danecek et al., 2021) was used to simulate 4 million paired-end 150 bp reads from each reference transcriptome using the following settings: -1 150 -2 150 -d 400 -N 4000000 -s 25 -e 0 -r 0 -R 0 -X 0. Simulated reads were aligned to the genome of a related species using the short read aligner BWA-MEM (Li & Durbin, 2009) with default parameters. Uniquely aligned and properly paired reads were selected and used to define the regions of contiguity between the transcriptome and the genome on which random sites were selected.

#### Selection of known target sites for probe design

For four species (*A. alba, P. halepensis, Pinus nigra*, and *T. baccata*), known target SNPs were selected from SPET panels developed within the European H2020 GENTREE project (https://www.gentree-h2020.eu/) to capture species-specific regions of interest, also including orthologous genes involved in stress responses and local adaptation. For *B. pendula, F. sylvatica, P. abies* and *Q. robur*, known target SNPs were selected from a dataset of genetic polymorphisms previously identified by an exome-capture sequencing experiment targeting orthologous genes involved in putative functions of interest such as response to stress, immune response, circadian clock and detection of abiotic stimulus (Milesi et al., 2024). For *Q. robur*, the known target set also included species-discriminatory markers (Guichoux et al., 2013; Nocchi et al., 2022; Reutimann et al., 2020) and additional candidate genes associated with responses to abiotic stressors (Homolka et al., 2013; Rellstab et al., 2016; Saleh et al., 2022; Trudić et al., 2021), pathogen resistance (Bartholomé et al., 2020), and phenology (Derory et al., 2009). For *F. excelsior*, target SNPs were selected from the multispecies ‘4TREE’ SNP chip (Archambeau et al., 2023; Guilbaud et al., 2020) produced within the framework of the European H2020 B4EST project (https://b4est.eu), comprising SNPs located in candidate genes associated with ash dieback disease resistance, emerald ash borer susceptibility, and self-incompatibility (Kelly et al., 2019; Sollars et al., 2017; Stocks et al., 2019). For *C. sativa*, known target positions included SNPs for varietal identification in European chestnuts (Nunziata et al., 2020), for species, hybrids and backcross characterization (Larue et al., 2021) and associated to chestnut gall wasp resistance (Gaudet et al., 2024). *M. sylvestris* known target positions were selected from a 20K genotyping array (Bianco et al., 2014), while *P. avium* targets were selected from a 6K genotyping array (Peace et al., 2012) and from markers related to cherry domestication (Pinosio et al., 2020).

### DNA extraction

For all species, DNA was extracted from 20-30 mg of dried leaves/needles per individual using NucleoSpin® Plant II kit (Macherey-Nagel). To reach the DNA quality and quantity required both for analyses by SPET (at least 500 ng of DNA/sample with a fluorimetric concentration of 20-50 ng/µl and with absorbance ratios of A_260_/A_280_ ≥ 1.7 and 1.6 ≤ A_260_/A_230_ ≤ 2.2) and SNP chip arrays (a fluorimetric DNA concentration of at least 30 ng/µl with absorbance ratios of 1.8 ≤ A_260_/A_280_ ≤ 2.0 and A_260_/A_230_ > 1.5), the manufacturer’s standard protocol was modified by adding 1ng/µl Proteinase K to the lysis buffer, increasing the lysis process up to 1 hour, and, only for *Quercus* spp., *P. avium* and *B. pendula*, by doubling the number of membrane washing cycles with the washing buffer. The CTAB-based lysis buffer was used for all species except *A. alba, P. halepensis, Populus alba* and *Populus nigra*, for which the SDS-based lysis buffer guaranteed higher quality and concentration of DNA.

### SNP array genotyping

Five species (*Pinus pinaster, Pinus pinea, P. sylvestris, Populus nigra* and *P. alba*) were genotyped at Thermo Fisher (Santa Clara, CA) using two different Axiom arrays developed within the European H2020 B4EST project. Specifically, *P. sylvestris* was genotyped using the PiSy50k Axiom array (Kastally et al., 2022) comprising a total of 47,712 SNPs, while the remaining species were genotyped using the multispecies 4TREE Axiom array (Archambeau et al., 2023; Guilbaud et al., 2020). This array comprises a total of 45,893 SNPs, of which 13,408 are for *Populus trichocarpa*, 13,407 are for *P. pinaster*, 5,671 are for *P. pinea* and the remaining 13,407 are for *F. excelsior*. The raw data for all species were obtained from Thermo Fisher and analyzed using the Axiom Analysis Suite software v5.2, by applying different sample quality and SNP quality thresholds for the two genera (*Populus*: DQC ≥ 0.82, QC call_rate ≥ 95, cr-cutoff ≥ 97, fld-cutoff ≥ 3.6, het-so-cutoff ≥ -0.1; *Pinus*: DQC ≥ 0.4, QC call_rate ≥ 85, cr-cutoff ≥ 85, fld-cutoff ≥ 3.2, het-so-cutoff ≥ -0.3).

### Library preparation, SPET sequencing and SNP detection

Genomic DNA was quantified using the Qubit 2.0 Fluorometer (Invitrogen, Carlsbad, CA). Libraries were prepared using the *Allegro Targeted Genotyping* protocol from Tecan Genomics (Männedorf, Zurich, Switzerland), using 100 ng of DNA as input and following the manufacturer’s instructions, including a 6 bp Unique Molecular Identifier (UMI) incorporated on each original DNA fragment. Libraries were quantified using the Qubit 2.0 Fluorometer, and their size was checked using the High Sensitivity DNA assay from Bioanalyzer (Agilent technologies, Santa Clara, CA). Sequencing was performed at IGA Technology Services (IGATech, Udine, Italy) facilities using a NovaSeq 6000 System (Illumina, San Diego, CA, USA) in paired-end mode (2×150 bp). BCL files from the instruments were processed using the manufacturer’s pipeline software to generate FASTQ sequence files.

Adaptor sequences and low quality 3’ ends were removed from DNA short reads using cutadapt (Martin, 2011) and ERNE-FILTER (Fabbro et al., 2013), with default parameters. After trimming, reads longer than 50 bp were aligned to the respective reference sequences (Table 1) using the short-read aligner BWA-MEM with default parameters (Li & Durbin, 2009). After alignment, duplicated sequences were removed using custom Perl script that leverages the UMIs added during library preparation (the script is available at 10.6084/m9.figshare.29560976). SNP calling was performed on uniquely aligned reads using the software package GATK version v4.5.0.0 (McKenna et al., 2010). First, *HaplotypeCaller* was run in GVCF mode to call potential variant sites at single-sample level in the SPET target regions. Then, the joint genotyping on the entire cohort of samples of each species was performed using *GenomicsDBImport* and *GenotypeGVCFs* tools.

### SNP filtering and population genetic statistics

From the gVCFs output by GATK, SNPs and invariant sites were temporarily separated for different filtering workflows. SNPs were isolated and filtered with GATK using *SelectVariants* --select-type-to-include SNP -- ignore-non-ref-in-types and *VariantFiltration* --filter-expression “QD < 2.0 || MQ < 40.0 || MQRankSum <-12.5”. SNPs resulting from ambiguous mapping between paralogous regions were removed based on elevated heterozygosity and read ratios using the HDplot method with options Hmax=0.6, RAFmin=0.2, RAFmax=0.8, Dmin=-10, Dmax=10 (McKinney et al., 2017). Genotype filtering was conducted with bcftools version 1.21 (Danecek et al., 2021). Only biallelic SNPs were retained, and only loci where “DP >= 6”, “GQ >= 20”, at least half the samples had a depth greater than six reads, and the minor allele was covered by at least three reads. For each dataset, samples with over 80% missing data were removed. Invariant sites were isolated from the original gVCF with *bcftools view* “--max-af 0 -V snps,indels,mnps,other” and filtered by “DP >= 6” where at least half the samples had a depth greater than six reads. Filtered invariant sites were combined with the respective filtered SNPs to create VCFs labelled version “V0”. Code used for filtering is available at DOI: 10.5281/zenodo.15820966.

The V0 filtered VCFs were used to identify and remove outlier and duplicate samples that may have originated while sampling or from other handling errors. To detect possible outliers, a principal component analysis (PCA) was conducted using the R package *snpR* (Hemstrom & Jones, 2023) with default settings. This information was combined with inference of individual admixture coefficients using sparse Non-Negative Matrix Factorization algorithms implemented in the *sNMF* function from R package *LEA* (Frichot et al., 2014) and interpreted in the light of the species biology known from literature and expert consultation. To explore species relatedness and identify putative duplicates, VCFs were formatted with PLINK version 1.9 (Chang et al., 2015) and the KING toolkit version 2.3.2 (Manichaikul et al., 2010) was run three times using the --related, --kinship, and --duplicates flags. If potential duplicates were likely to be natural clones, they were retained; otherwise, one of each pair was removed to produce V1 VCFs. Outliers that were clearly the result of sample mishandling were also removed from V1 files.

Expected heterozygosity (*H*_S_) and Weir & Cockerham’s *F*_ST_ (Weir & Cockerham, 1984) were estimated at species level using R package *hierfstat* v0.5-11 (Goudet, 2005). Bootstrapping (1,000) was used to obtain a 95% confidence interval for the *F*_ST_ values. Nucleotide diversity was estimated using pixy version 1.2.7.beta1 (Korunes & Samuk, 2021) on V1 filtered VCF files.

To identify SNPs that discriminate among *Quercus robur, Q. petraea*, and *Q. pubescens*, raw VCF files from individuals of each species were merged using bcftools. SNPs were then extracted and filtered using GATK’s SelectVariants and VariantFiltration tools, as previously described. Additional filtering was performed with bcftools to retain only biallelic SNPs with a minimum depth ≥ 6, genotype quality ≥ 20, and present in at least 80% of individuals. A PCA was conducted to confirm species identity and exclude putative hybrids or misassigned individuals. For each species, a filtered VCF including only confirmed individuals was generated, and allele frequencies were computed using vcftools. SNPs showing large allele frequency differences (≥ 0.9) between at least two of the three species were retained as species-discriminant loci.

## Results

### SPET panels performance

The average number of sequenced read pairs per sample ranged from about 1.3 million in *P. avium, Q. ilex*, and *Quercus suber* to about 3.4 million in *Pinus nigra* (Table 2). The alignment rate to the whole genome, considering only uniquely mapped reads, was highly variable among species, ranging from 32.5% when aligning against the highly fragmented 10 Gb genome of *P. abies* to 91.2% when aligning to the pseudomolecule-level 800 Mb genome of *F. excelsior*. In each species, the majority of the 10K designed probes were effective, producing an average sequence coverage of at least 6X at the target site in at least half of the genotyped individuals. However, for a group of related species belonging to the Fagaceae family and genotyped with the FORGENIUS-MultiSp50K SPET panel (*C. sativa, F. sylvatica*, and *Quercus* spp.), the number of effective probes exceeded 10K due to the cross-species transferability of a subset of probes. For each of these related species, Table S3 shows the number of effective probes, categorized according to the species they were originally designed for. The average depth obtained at the target regions captured by the working probes ranged from 18.3X in *A. nebrodensis* to 85.1X in *F. excelsior*. The size of the captured genome (i.e., positions of the genome having a sequence coverage ≥6x in at least half of the genotyped samples) ranged from 1.12 Mb in *P. abies* to 3.78 Mb in *F. excelsior*.

SPET panels demonstrated high performance when applied to genotype related species. For instance, nearly 9,300 of the 10,000 probes designed for *A. alba* successfully genotyped *A. nebrodensis* samples. Similar results were observed in most of the *Quercus* species, where 10,000 probes were designed based on the *Q. robur* genome and 3,500 probes based on the *Q. ilex* genome. Of these, 10,700, 11,000, and 4,100 probes were effective in genotyping *Q. pubescens, Q. petraea*, and *Q. suber*, respectively. Additionally, we tested the SPET panel of *P. halepensis* on a small dataset of *Pinus heldreichii* samples (not included in the 23 species selected in FORGENIUS), and approximately 6,700 probes were effective. To visually inspect the genomic distribution of the SPET panels, we plotted the distribution of the working probes in 250 Kb windows for all species with a reference genome at the pseudomolecule level (Figure 2 and Supplementary Figures S1-S9). Probes were generally well distributed, although consecutive windows without probes were more frequent in species for which probes were designed on the transcriptome (*C. sativa* and *T. glaberrima*) and in species with large genomes (*P. halepensis, Pinus nigra* and *T. baccata*). As an example, Figure 2 shows the distribution of *Q. robur* probes mapped to its own 0.8 Gb genome (top) and *Pinus nigra* probes mapped to the 25.6 Gb *Pinus tabuliformis* genome (bottom). Despite the expected presence of windows without probes in the *P. tabuliformis* genome, the distribution remains relatively uniform across its 12 chromosomes.

**Figure 2.**
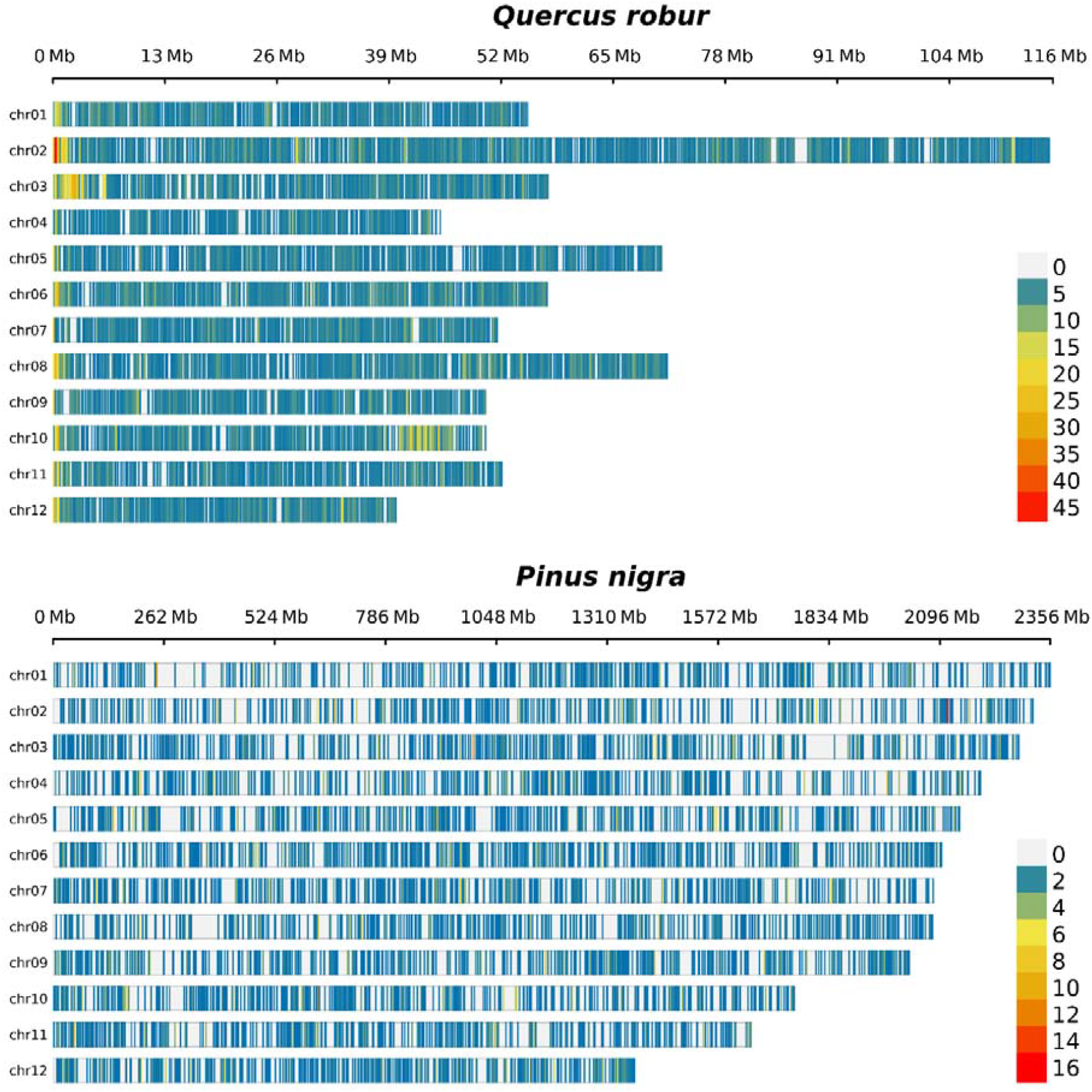
Distribution of Quercus robur probes with respect to the Q. robur PM1N genome (top) and Pinus nigra probes on the Pinus tabuliformis genome (bottom), represented as number of probes in 250-Kb windows.

### SNP detection and genomic characterization

We genotyped 300 Genetic Conservation Units (GCUs) across 23 forest tree species using either the SPET technology (Scaglione et al., 2019) or the Axiom array-based genotyping technology (Thermo Fisher, Santa Clara, CA). For the species genotyped using SPET technology, we obtained tens of thousands of SNPs, whereas for those genotyped with Axiom arrays, the number of SNPs was more limited and proportional to the number of markers included in each array (Table 3). The species with the lowest number of detected SNPs was *P. alba* (3,031), where most of the markers resulted monomorphic in the 126 genotyped samples, reflecting the use of an array designed for Populus nigra. Among the species genotyped by SPET, the lowest number of SNPs was detected in A. nebrodensis (16,828), a narrow-endemic species for which the only 30 existing individuals were genotyped. To assess the reproducibility of SPET genotyping, a total of 48 samples (at least one per species, except for *A. nebrodensis*) were genotyped twice. On average, 99.1% of genotype calls were concordant between replicates (Figure S10A). Among the discordant calls, the majority (80.2%) were homozygous reference in one replicate and heterozygous in the other, followed by calls that were heterozygous in one replicate and homozygous alternative in the other (16.2%). The least frequent discordances (3.6%) occurred when one replicate was homozygous reference and the other homozygous alternative. Overall, the mean depth at concordant calls was higher compared to discordant ones (Figure S10B). Finally, SNPs in the V1 VCF files were used to calculate, at the species level, expected heterozygosity (*H*_S_) and Weir & Cockerham’s *F*_ST_, as well as nucleotide diversity only for SPET-genotyped species. The results are reported in Table 3.

**Table 3.** Number of successfully genotyped samples, name of the genotyping tool, number of SNPs characterized, as well as expected heterozygosity (*H*_S_), nucleotide diversity (*π*) and genetic differentiation (*F*_ST_) per species.

| Species | Genotyped samples | Genotyping tool | Number of SNPs | $H_s$ | $\pi$ | $F_{ST}$ |
| --- | --- | --- | --- | --- | --- | --- |
| *Abies alba* | 413 | FORGENIUS-Aalb10K | 66,940 | 0.084 | 0.00201 | 0.107 |
| *Abies nebrodensis* | 24 | FORGENIUS-Aalb10K | 16,828 | 0.243 | 0.00265 | NA |
| *Betula pendula* | 351 | FORGENIUS-MultiSp60K | 167,558 | 0.061 | 0.00387 | 0.109 |
| *Castanea sativa* | 312 | FORGENIUS-MultiSp50K | 64,212 | 0.150 | 0.00231 | 0.236 |
| *Fagus sylvatica* | 370 | FORGENIUS-MultiSp50K | 113,379 | 0.098 | 0.00352 | 0.116 |
| *Fraxinus excelsior* | 331 | FORGENIUS-MultiSp60K | 259,272 | 0.082 | 0.00538 | 0.163 |
| *Malus sylvestris* | 347 | FORGENIUS-MultiSp60K | 114,942 | 0.115 | 0.00411 | 0.087 |
| *Picea abies* | 396 | FORGENIUS-MultiSp50K | 72,553 | 0.066 | 0.00370 | 0.075 |
| *Pinus halepensis* | 344 | FORGENIUS-Phal10K | 34,671 | 0.080 | 0.00148 | 0.194 |
| *Pinus nigra* | 369 | FORGENIUS-MultiSp60K | 93,975 | 0.061 | 0.00339 | 0.078 |
| *Pinus pinaster* | 367 | 4TREE Axiom array | 9,418 | 0.266 | NA | 0.125 |
| *Pinus pinea* | 144 | 4TREE Axiom array | 4,105 | 0.171 | NA | 0.442 |
| *Pinus sylvestris* | 368 | Pisy50k Axiom array | 45,998 | 0.299 | NA | 0.058 |
| *Populus alba* | 126 | 4TREE Axiom array | 3,031 | 0.095 | NA | 0.072 |
| *Populus nigra* | 329 | 4TREE Axiom array | 10,255 | 0.207 | NA | 0.183 |
| *Prunus avium* | 262 | FORGENIUS-MultiSp60K | 85,249 | 0.084 | 0.00181 | 0.207 |
| *Quercus ilex* | 320 | FORGENIUS-MultiSp50K | 217,986 | 0.053 | 0.00513 | 0.076 |
| *Quercus petraea* | 301 | FORGENIUS-MultiSp50K | 244,822 | 0.050 | 0.00407 | 0.048 |
| *Quercus pubescens* | 121 | FORGENIUS-MultiSp50K | 170,451 | 0.066 | 0.00443 | 0.013 |
| *Quercus robur* | 387 | FORGENIUS-MultiSp50K | 175,185 | 0.072 | 0.00392 | 0.046 |
| *Quercus suber* | 385 | FORGENIUS-MultiSp50K | 71,467 | 0.106 | 0.00260 | 0.054 |
| *Taxus baccata* | 387 | FORGENIUS-MultiSp60K | 52,533 | 0.176 | 0.00306 | 0.196 |
| *Torminalis glaberrima* | 244 | FORGENIUS-MultiSp50K | 51,350 | 0.142 | 0.00237 | 0.111 |

For the subset of species within the Fagaceae family where cross-species transferability of some probes was observed, we assessed whether the use of sequences captured by probes from different species introduced any biases in the estimation of genetic diversity. We calculated observed heterozygosity and minor allele frequency (MAF), stratifying the data according to the origin of the probes. Our analysis revealed no significant differences between the results obtained from species-specific probes and those captured by probes from other species, suggesting that cross-species probe transfer did not affect the genetic diversity estimates (Figure S11).

For *M. sylvestris* and *F. excelsior* a portion of the known target sites included in the SPET panels were derived from two previously developed SNP arrays: an Illumina Infinium array targeting 20K SNPs for *M. sylvestris* (Bianco et al., 2014) and the multispecies ‘4TREE’ Axiom SNP chip for *F. excelsior* (Guilbaud et al., 2020). This allowed us to compare the genetic diversity estimates obtained for these two species using the two different technologies, either by considering only the subset of SNPs included in the arrays (“array SNPs”) or by analyzing *de novo* SNPs identified in regions captured by random probes (“random SNPs”). In both species, we observed a higher MAF and, therefore, observed heterozygosity, in the array SNP set compared to the *de novo* SNP set (Figure 3), as expected due to the preference of SNPs at intermediate frequencies in SNP selection for arrays. This effect was more pronounced in *F. excelsior*, where the differences in the distributions of MAF and heterozygosity were both statistically significant (one-sided Wilcoxon rank-sum test p-values 2.2e-16 and 1.7e-6, respectively). In contrast, in *M. sylvestris*, only the difference in the MAF distribution was statistically significant (p-value 2.2e-16). The overestimation of diversity metrics observed in array SNPs determined an underestimation of *F*_ST_ ranging from 5% in *F. excelsior* to 20% in *M. sylvestris*. When the same statistics were calculated for the entire dataset, the estimates were comparable to those obtained from the random probes, indicating that any potential bias introduced by the known target SNPs is mitigated when the data are combined. This is inherently linked to the much larger number of random than target SNPs per probe usually obtained by SPET genotyping (Scaglione et al., 2019).

**Figure 3.**
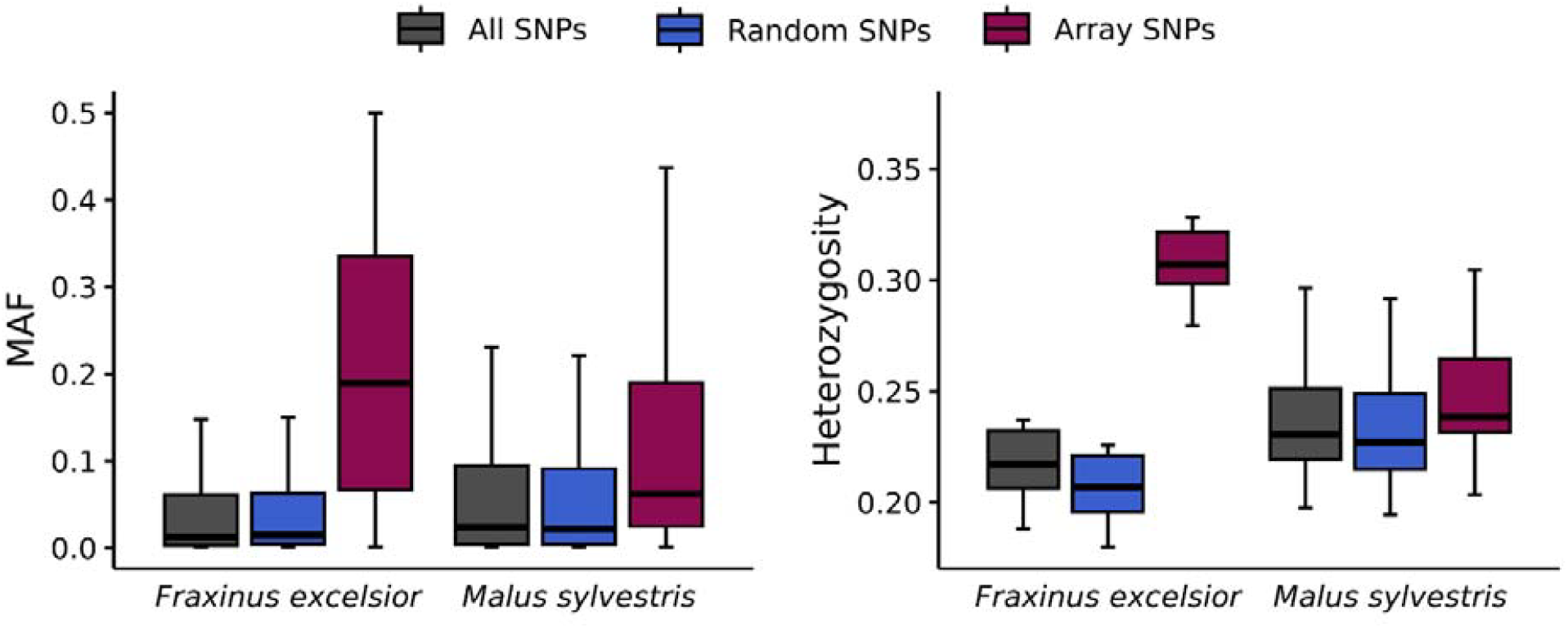
Comparison of genetic diversity estimates based on SPET and SNP array technologies. Boxplots show the distribution of MAF (left panel) and observed heterozygosity (right panel) in *F. excelsior* and *M. sylvestris*, using either the entire dataset (All SNPs), *de novo* SNPs identified in regions captured by random probes (Random SNPs), or SNPs included in the SNP arrays (Array SNPs). Outliers in the boxplots are not displayed to enhance readability.

The large number of new markers discovered by SPET genotyping enables multi-purpose applications beyond the original goal of improving the characterization of GCUs for future genetic monitoring. For instance, the SPET panel developed for *Q. robur* has the potential to capture approximately 200 markers that have been reported to have high discriminatory power among different white oak species (Guichoux et al., 2013; Kremer et al., 2024; Reutimann et al., 2020), listed in Table S4. We also searched for a new set of diagnostic markers by selecting genomic positions where the difference in the frequency of the alternative allele between at least two of the three white oak species was greater than 0.9. Through this approach, we identified a total of 196 new markers with high discriminatory power, as listed in Table S5.

## Discussion

In the framework of the H2020 European project FORGENIUS, new genomic resources were developed and data produced for 23 forest tree species spanning from rare and scattered (e.g., *A. nebrodensis* and *T. glaberrima*) to widespread, economically relevant ones (e.g., *F. sylvatica, P. abies* and *P. sylvestris*). By designing ad-hoc SPET panels and employing already available SNP array chips, we identified a total of 1,852,293 SNPs across 7,192 trees, covering about 50 Mb of genomic sequence. SPET panels were specifically designed to obtain a wide plateau of genomic information, while minimizing potential biases affecting estimates, for the genetic monitoring and the assessment of possible maladaptation under future environmental conditions across the European network of forest Genetic Conservation Units (GCUs). The main features and the potential for a wider usage of the resources presented here are discussed in the following.

### Performance and key features of FORGENIUS SPET panels

SPET technology allowed us to generate an extensive set of genetic markers across the analyzed species, regardless of genome size or complexity. In fact, even if a higher average coverage was generally obtained in species with smaller and less repetitive genomes, the newly developed SPET designs consistently provided a valuable set of genetic markers well-distributed across the genomes for all analyzed species, including those with larger and more repetitive genomes. Notably, the SPET probes demonstrated a high transferability rate among related species, as exemplified by the successful genotyping of *A. nebrodensis* using probes designed for *A. alba* and the effective cross-species performance observed in the *Quercus* genus. High transferability represents a significant advantage compared to other methods, such as SNP-genotyping arrays, where inter-species transferability is often limited (Geraldes et al., 2013; Peace et al., 2012). For example, in the case of *P. alba*, we found a very limited number of polymorphic markers when using the 4TREE array developed for *Populus nigra*. This cross-species transferability was further supported by results obtained in species of the Fagaceae family genotyped using the Forgenius-MultiSp50K SPET panel (*C. sativa, F. sylvatica*, and *Quercus* spp.), where the number of effective probes exceeded 10,000 due to the successful amplification of a subset of probes across species. Importantly, the high concordance of genetic diversity estimates obtained using either species-specific probes or probes originally designed for related species suggests that any ascertainment bias potentially introduced by cross-species probe application is minimal, if present at all. This versatility suggests that the SPET panels developed in this study could serve as valuable tools for studying genetic diversity in other tree species beyond those included in this work (e.g., the nine Mediterranean firs other than *A. nebrodensis*, Alizoti et al., 2011; the red-listed *Picea omorika*, Mataruga et al., 2020) Additionally, the ability to employ these panels for comparative analyses across species highlights their utility in evolutionary and ecological studies, offering a scalable and robust approach to genomic characterization across diverse taxa. Moreover, the availability of multispecies and cross-transferable SPET panels can help overcome scalability issues often encountered in genotyping projects, particularly when only a limited number of samples need to be genotyped per species. The ability to process multiple species simultaneously in a single run may reduce per-sample costs and increase genotyping efficiency.

The high transferability of SPET probes among related species highlights the potential of SPET genotyping not only for broad genetic characterization but also for the identification of species differentiation markers. In the case of white oaks, the SPET panel including *Quercus* species (FORGENIUS-MultiSp50K) successfully captured approximately 200 markers previously reported as useful for species differentiation (Guichoux et al., 2013; Kremer et al., 2024; Reutimann et al., 2020) and allowed us identify additional 196 markers with high resolution power in distinguishing species. These newly discovered markers, based on range-wide discovery panels, significantly enhance the ability to differentiate between closely related white oak species, confirming the potential of SPET panels as a powerful tool for species delimitation. Among the wide range of other possible applications, the FORGENIUS SPET panels could also prove useful in reconstructing the origin of material in forest plantations and breeding programs, which is attracting attention for its straightforward applications in predicting the performance of different provenances in relevant climatic setups (e.g., Milesi et al., 2019; Oggioni et al., 2024). The ability to both genotype known markers and discover new ones highlights the effectiveness of SPET genotyping in evolutionary and taxonomic studies (Barchi et al., 2019; Gramazio et al., 2020), providing a scalable and transferable framework for marker discovery across a wide range of species.

In *F. excelsior* and *M. sylvestris*, we were able to evaluate the ability of the SPET designs to capture genetic diversity compared to previously developed SNP arrays showing that SPET panels specifically tailored for these species represent a significant advancement in accurately assessing their range-wide genetic diversity. Unlike SNP arrays, which are prone to ascertainment bias due to the selection of target polymorphisms from specific populations or studies, the SPET design allowed for the *de novo* detection of SNPs across the entire genome. The differences in diversity and differentiation metrics observed between array SNPs and random SNPs highlight the impact of ascertainment bias in pre-designed arrays, both in *F. excelsior*, where the disparity was statistically significant for both diversity metrics, and in M. sylvestris where the effect was more marked on *F*_ST_ underestimation. Also considering that SPET data, including information on invariant sites, permits estimating an important additional diversity metric, i.e. nucleotide diversity, they overall enable a more comprehensive characterization of GCUs’ genetic composition and more robust conclusions about their evolutionary history and adaptive potential.

### Applications of FORGENIUS genomic resources to the conservation of European forests

The European network of GCUs was established almost 20 years ago by the EUFGIS project (http://www.eufgis.org) and has never been systematically characterized for its genetic features. To this aim, it is essential to have tools available for gathering comparable information for a large number of species, which was one of the main goals of the ongoing EU H2020 FORGENIUS project. Genomic resources, both in terms of genotyping tools and data, are in fact basic requirements for starting a genetic monitoring program aimed at assessing GCUs’ adaptive capacity, a crucial component of populations’ vulnerability (IPCC, 2007), through time. Genetic monitoring, i.e. the continuous assessment of populations’ genetic diversity, is advocated as a key action to properly conserve and manage natural genetic resources (Pearman et al., 2024) and comply with international commitments for the conservation of biological diversity (e.g., the Convention on Biological Diversity, 2022).

The genomic resources presented here proved to be an efficient and affordable tool to characterize the genetic diversity of the European network of GCUs, while providing insights on both future adaptability and past demography, which are key aspects to interpret current forest dynamics and orient conservation strategies. Our parameter estimations at the species level shows the large variety of past demographic histories experienced by FORGENIUS focal species, from the highly panmictic white oaks and *P. sylvestris* (*F*_ST_ □0.05 calculated on populations encompassing a large part of their natural distribution) to highly structured ones reflecting historical fragmentation (e.g., *T. baccata*, Mayol et al., 2015), and/or factors related to limited dispersal and human exploitation (e.g., *C. sativa*, Mattioni et al., 2017; *P. avium*, Pinosio et al., 2020; *P. pinea*, Jaramillo-Correa et al., 2020). Our estimates of nucleotide diversity and expected heterozygosity are usually in line with what was found in previous literature, confirming general trends already highlighted in multi-species studies of forest trees (e.g., the high intraspecific diversity of white oaks as compared to other European anemophilous forest tree species, Milesi et al., 2024), with novel insights on less studied species, such as the relatively high nucleotide diversity of the critically endangered *A. nebrodensis* unique population. Among the FORGENIUS focal species, *M. sylvestris* is the progenitor of a domesticated species, and *P. avium* of widely used cultivars. In both cases, the nucleotide diversity detected in GCUs is relatively similar to estimates obtained studying domesticated varieties (Micheletti et al., 2011; Pinosio et al., 2020). This suggests potentially critical issues in the selection and distribution of GCUs, which would be expected to have higher portion of genetic diversity than related, domesticated varieties. Finally, the presented datasets will generate what, to our knowledge, are the first estimates of nucleotide diversity for species that require special conservation actions (e.g., *A. nebrodensis*, del Valle et al., 2024) and/or will be fundamental elements in future reforestation programs to increase species diversity of European forests (*T. glaberrima*, Afifi et al., 2025).

Although specific analysis at the GCU level for each species will be the subject of forthcoming studies, for the scientific community interested in using the presented datasets, it is worth noting that the more elaborated VCF files provided (V1) still contain, for instance, hybrid individuals and trees potentially belonging to selected cultivars, because these individuals are considered highly informative on the status of single GCUs as part of the diversity that is preserved within them. These individuals could be of course removed in specific data analyses as needed.

### Final remarks and future directions

The FORGENIUS SPET panels and data presented here are intended to describe the status of the European network of forest GCUs, thus representing a first step in developing monitoring programs for the forest genetic resources conserved in this infrastructure. Data are available for 23 forest tree species and, considering the high transferability observed in this study at the genus and even family level, the same tools could be applied to a much larger number of species. In addition, our work provides the basis to design SPET panels with promising features on other species as the necessary resources (e.g., full transcriptome and genome reference sequences) become available. For species with limited genome size (ca. 1 Gbp), it will be possible to switch to whole-genome sequencing soon, although its applicability to thousands of samples is still economically unsuitable. However, even in the case of a strong acceleration on this front, the data produced using the FORGENIUS approach could easily be merged with deeper genomic analyses. For species with extra-long and complex genomes, such as conifers, the tools presented here could represent a convenient option for characterizing the range-wide genetic diversity of hundreds of populations, which is the sample size required by comprehensive genetic monitoring programs.

The possibility of using a mixture of genetic markers in candidate genes as well as randomly distributed throughout the genome makes SPET data particularly flexible to cover a wide range of research aims, from those related to Gene-Environment Association (GEA) analyses (e.g., genomic offset estimation, Rellstab & Keller, 2025) to the assessment of genetic load, mating patterns and demographic analyses, or more conservation-oriented goals such as those of Spatial Conservation Planning (Vajana et al., 2024). The FORGENIUS SPET panels have been proved ideal tools also to deepen issues related to hybridization and species delimitations, with potentially high accuracy at the whole distribution level, which was problematic when diagnostic markers were developed on geographically limited discovery panels (e.g., Kremer et al. 2024). Finally, we showed how SPET panels transferred to congeneric species could be particularly suitable for comparative genomics analyses, thus providing essential information to forecast forest trees responses to environmental fluctuations (Milesi et al., 2024; Yeaman et al., 2016).

### Data Accessibility Statement

Raw and filtered genotyping data have been uploaded either to the NCBI Sequence Read Archive or the Figshare platform and will be released upon acceptance of the paper.

Raw FASTQ data are available on the Sequence Read Archive (SRA) on NCBI under the BioProject IDs PRJNA1183629 (*Abies alba*), PRJNA1187359 (*Abies nebrodensis*), PRJNA1196663 (*Betula pendula*), PRJNA1199760 (*Castanea sativa*), PRJNA1245097 (*Fagus sylvatica*), PRJNA1198695 (*Fraxinus excelsior*), PRJNA1202045 (*Malus sylvestris*), PRJNA1245134 (*Picea abies*), PRJNA1187483 (*Pinus halepensis*), PRJNA1191837 (*Pinus nigra*), PRJNA1214746 (*Prunus avium*), PRJNA1246490 (*Quercus ilex*), PRJNA1247253 (*Quercus petraea*), PRJNA1216914 (*Quercus pubescens*), PRJNA1247390 (*Quercus robur*), PRJNA1247367 (*Quercus suber*), PRJNA1207918 (*Taxus baccata*), PRJNA1247414 (*Torminalis glaberrima*). Axiom SNP array raw data for *Pinus pinaster, Pinus pinea, Pinus sylvestris, Populus alba* and *Populus* nigra are available on Figshare at 10.6084/m9.figshare.28787597. Filtered VCF for all species are available on Figshare at 10.6084/m9.figshare.28741400.

### FORGENIUS Consortium Co-authors

Marc Busuldu, Vega Garcia-Segura, Ana Hernández-Serrano, Sonia Hernando, Maurizio Mencuccini, Joan Prunera-Olivé, Carla Rodrigo-González, Adolfo Sanmartín-Arévalo, Martí Solé-Xanxo, Laura Wynne Stewart (CREAF, UAB Campus Bellaterra, Edifici C, 08193 Cerdanyola del Vallès Barcelona, Spain); Ana Cabanillas (Department of Environment and Tourism, Government of Aragón, Spain); Jonathan Feichter, Berthold Heinze Department of Forest Growth, Silviculture and Genetics; Austrian Research Centre for Forests, BFW, Seckendorff-Gudent-Weg 8, A-1131 Vienna, Austria); Akkin Semerci (Department of Forestry and Forest Products, Niksar Vocational School, Tokat Gaziosmanpaşa University, Tokat, Türkiye); Katrin Heer, Jill Sekely (Eva Mayr-Stihl Professorship of Forest Genetics, Faculty of Environment and Natural Resouces, Albert-Ludwigs Universität Freiburg, Bertoldstraße 17, 79111 Freiburg, Germany); Stuart A’Hara, Joan Cottrell (Forest Research, Northern Research Station, Roslin, Midlothian, EH25 9SY, UK); Eduardo Notivol (Government of Aragon Avda. Montañana 930, 50059 Zaragoza, Spain); Christophe Gauvrit, Marie-Gabrielle Harribey, Joan Hochet, Baptiste Laffitte, Matteo Nasuti, Annie Raffin (INRAE, UEFP, 69 route d’Arcachon, 33612 Cestas Cedex, France); Thomas Francisco, Adélaïde Theraroz (INRAE, Univ. Bordeaux, BIOGECO, 69 route d’Arcachon, 33612 Cestas Cedex, France); Pierre-Jean Dumas, Florence Jean, Nicolas Mariotte (INRAE, URFM, 228 Route de l’Aérodrome, 84914 Avignon Cedex 9); Maurizio Marchi (Institute of Biosciences and BioResources - IBBR, National Research Council - CNR, Via Madonna del Piano 10, 50019 Sesto Fiorentino, Firenze, Italy); Ricardo Alía, Francisco J. Auñón, Diana Barba, Maria Regina Chambel, Fernando del Caño, María Carmen García Barriga, José Manuel García del Barrio, Delphine Grivet, Rodrigo Pulido Sanz (Instituto de Ciencias Forestales ICIFOR-INIA, Consejo Superior de Investigaciones Cientificas CSIC, Crta. A Coruña Km7.5, 28040 Madrid, Spain); Ana Iordan, Flaviu Popescu, Dragos Postolache, Daniel Suciu (National Institute for Research and Development in Forestry “Marin Dracea”, Bv. Eroilor nr.128, Voluntari, Ilfov, Romania); Leena Yrjänä (Natural Resources Institute Finland Luke, Latokartanonkaari 9, 00790 Helsinki, Finland); Egbert Beuker, Henri Hämäläinen, Hanna-Riikka Haurinen (Natural Resources Institute Finland Luke, Vipusenkuja 5, 57200 Savonlinna, Finland); Christian Mestre-Runge, Lars Opgenoorth, Christian Reudenbach (Plant Ecology and Geobotany, Philipps Universität Marburg, Karl-von-Frisch-Straße 8, 35043 Marburg, Germany); Marko Bajc, Gregor Božič, Rok Damjanić, Natalija Dovč, Luka Krajnc, Marija Kravanja, Gal Oblišar, Boris Rantaša, Gregor Skoberne, Barbara Štupar, Urša Vilhar, Marjana Westergren (Slovenian Forestry Institute, Večna pot 2, 1000 Ljubljana, Slovenia); Stephen Cavers, Annika Perry (UK Centre for Ecology & Hydrology, Bush Estate, Penicuik, UK. EH31 2HY).

## Author Contributions

S Pinosio, F Bagnoli, C Avanzi, I Spanu, SC González-Martínez, T Pyhäjärvi, I Scotti, GG Vendramin and A Piotti designed the research. F Bagnoli, C Avanzi, I Spanu, MB Castellani and A Frascella performed the lab work. S Pinosio, SL McEvoy and S Olsson analysed the data. A Piotti, T Pyhäjärvi, M Westergren, I Scotti, GG Vendramin, D Grivet, R Alia, SC González-Martínez, S Cavers, B Heinze and C Avanzi supervised the selection of samples to create V1 datasets. F Bagnoli, C Avanzi, I Spanu, SC González-Martínez, MB Castellani, A Frascella, E Vajana, S Olsson, I Scotti, GG Vendramin, A Piotti and the FORGENIUS Consortium performed the field work. S Pinosio, A Piotti, SL McEvoy and S Olsson wrote the manuscript. F Bagnoli, C Avanzi, E Vajana, SC González-Martínez, T Pyhäjärvi, I Scotti and GG Vendramin critical revised the manuscript for important intellectual content. All Authors revised the final version of the manuscript.

## Acknowledgements

This work was funded by the European Union (EU)’s Horizon 2020 Research and Innovation Program under grant agreement No. 862221 (FORGENIUS) and took advantage of the resources available from the EU-funded projects under grant agreements No. 773383 (B4EST), No. 676876 (GENTREE), and No. 101107604 (MedForAct). The views and opinions expressed are, however, those of the authors only and do not necessarily reflect those of the European Union. Neither the European Union nor the Granting Authority can be held responsible for them. We would also like to thank the ITINERIS project - Piano Nazionale Ripresa e Resilienza (PNRR) Missione 4, Componente 2, Investimento 3.1 “Fondo per la realizzazione di un sistema integrato di infrastrutture di ricerca e innovazione” - Italian Integrated Environmental Research Infrastructures System, for supporting the implementation of CNR-IBBR laboratory infrastructure.

We wish to thank all the Parks and Reserves where sampling sites are located, as well as private owners, for permissions and logistic support. We are also very grateful to S Scalabrin and IGATech personnel, MJM Smulders, M Troggio, N Howard, L Bianco, M Gaudet, C Boggi, for their valuable contribution at various stages of the research, F Al-Awad for her contribution to the sampling of *Populus nigra* in the Austrian GCU AUT00284, and the Unité Expérimentale Forêt Pierroton’ (UEFP, INRAE; https://doi.org/10.15454/1.5483264699193726E12) for field assistance. We also thank H Wildhagen (HAWK Hochschule für angewandte Wissenschaft und Kunst Hildesheim/Holzminden/Göttingen, Germany) and O Gailing (Georg-August-Universität Göttingen, Germany) for sharing the 90K SPET panels of *Q. petraea* and *F. sylvatica*, developed within the DroughtMarkers project (funding provided by FNR-Waldklimafonds - 2218WK43A4 and 2218WK43B4), which we used to design the probes for *Q. robur* and *F. sylvatica*.

## References

Acquadro, A., Marinoni, D. T., Sartor, C., Dini, F., Macchio, M., & Botta, R. (2019). Transcriptome characterization and expression profiling in chestnut cultivars resistant or susceptible to the gall wasp Dryocosmus kuriphilus. Molecular Genetics and Genomics 2019 295:1, 295(1), 107–120. 10.1007/S00438-019-01607-2

Afifi, L., Szukala, A., Klumpp, R., Tremetsberger, K., & Konrad, H. (2025). Monitoring genetic diversity of Torminalis glaberrima for resilient forests in the face of population fragmentation. Annals of Botany, mcaf010. 10.1093/aob/mcaf010

Alizoti, P., Fady, B., Prada, M. A., & Vendramin, G. G. (2011). EUFORGEN Technical Guidelines for genetic conservation and use of Mediterranean firs (Abies spp.). Bioversity International, Rome, Italy, 6 p.

Aravanopoulos, F., Tollefsrud, M. M., Graudal, L., Koskela, J., Kätzel, R., Soto, A., Nagy, L., Pilipović, A., Zhelev, P., Božič, G., & Bozzano, M. (2015). Genetic monitoring methods for genetic conservation units of forest trees in Europe. Bioversity International, Rome, Italy, 62 p.

Archambeau, J., Bianchi, S., Buiteveld, J., Callejas-Díaz, M., Cavers, S., Hallingbäck, H., Kastally, C., de Miguel, M., Mutke, S., Sánchez, L., Whittet, R., González-Martínez, S. C., & Bastien, C. (2023). Managing forest genetic resources for an uncertain future: Findings and perspectives from an international conference. Tree Genetics & Genomes, 19(3), 26. 10.1007/s11295-023-01603-z

Barchi, L., Acquadro, A., Alonso, D., Aprea, G., Bassolino, L., Demurtas, O., Ferrante, P., Gramazio, P., Mini, P., Portis, E., Scaglione, D., Toppino, L., Vilanova, S., Díez, M. J., Rotino, G. L., Lanteri, S., Prohens, J., & Giuliano, G. (2019). Single Primer Enrichment Technology (SPET) for High-Throughput Genotyping in Tomato and Eggplant Germplasm. Frontiers in Plant Science, 10, 1005. 10.3389/fpls.2019.01005

Bartholomé, J., Brachi, B., Marçais, B., Mougou-Hamdane, A., Bodénès, C., Plomion, C., Robin, C., & Desprez-Loustau, M. L. (2020). The genetics of exapted resistance to two exotic pathogens in pedunculate oak. New Phytologist, 226(4), 1088–1103. 10.1111/NPH.16319

Bianco, L., Cestaro, A., Sargent, D. J., Banchi, E., Derdak, S., Guardo, M. D., Salvi, S., Jansen, J., Viola, R., Gut, I., Laurens, F., Chagné, D., Velasco, R., Weg, E. V. D., & Troggio, M. (2014). Development and Validation of a 20K Single Nucleotide Polymorphism (SNP) Whole Genome Genotyping Array for Apple (Malus × domestica Borkh). PLOS ONE, 9(10), e110377. 10.1371/JOURNAL.PONE.0110377

Bruxaux, J., Zhao, W., Hall, D., Curtu, A. L., Androsiuk, P., Drouzas, A. D., Gailing, O., Konrad, H., Sullivan, A. R., Semerikov, V., & Wang, X.-R. (2024). Scots pine – panmixia and the elusive signal of genetic adaptation. New Phytologist, 243(3), 1231–1246. 10.1111/nph.19563

Budde, K., Zormpa, O. G., Wilhelmi, S., Vucetic, B., Müller, M., Gailing, O., Ciocîrlan, M.-I.-C., Ciocîrlan, E., Curtu, A. L., Targem, M., & Wildhagen, H. (2024). Differences in fine-scale spatial genetic structure of European beech populations along elevational gradients. Research Square. 10.21203/rs.3.rs-4559673/v1

Chang, C. C., Chow, C. C., Tellier, L. C., Vattikuti, S., Purcell, S. M., & Lee, J. J. (2015). Second-generation PLINK: Rising to the challenge of larger and richer datasets. GigaScience, 4, 7. 10.1186/s13742-015-0047-8

Convention on Biological Diversity. (2022). First draft of the post-2020 global biodiversity framework. Conference of the parties to the Convention of Biological Diversity (Fifteenth meeting - Part II) (No. Agenda item 9A). Montreal, Canada.

Danecek, P., Bonfield, J. K., Liddle, J., Marshall, J., Ohan, V., Pollard, M. O., Whitwham, A., Keane, T., McCarthy, S. A., & Davies, R. M. (2021). Twelve years of SAMtools and BCFtools. GigaScience, 10(2), 1–4. 10.1093/GIGASCIENCE/GIAB008

de Vries, S. M. G., Alan, M., Bozzano, M., Burianek, V., Collin, E., Cottrell, J., Ivankovic, M., Kelleher, C. T., Koskela, J., Rotach, P., Vietto, L., & Yrjänä, L. (2015). Pan-European strategy for genetic conservation of forest trees and establishment of a core network of dynamic conservation units. European Forest Genetic Resources Programme (EUFORGEN), Bioversity International, Rome, Italy. xii + 40 p.

del Valle, J. C., Arista, M., Benítez-Benítez, C., Ortiz, P. L., Jiménez-López, F. J., Terrab, A., & Balao, F. (2024). Genomic-Guided Conservation Actions to Restore the Most Endangered Conifer in the Mediterranean Basin. Molecular Ecology, n/a(n/a), e17605. 10.1111/mec.17605

Derory, J., Scotti-Saintagne, C., Bertocchi, E., Dantec, L. L., Graignic, N., Jauffres, A., Casasoli, M., Chancerel, E., Bodénès, C., Alberto, F., & Kremer, A. (2009). Contrasting relations between diversity of candidate genes and variation of bud burst in natural and segregating populations of European oaks. Heredity 2010 105:4, 105(4), 401–411. 10.1038/hdy.2009.170

Exposito-Alonso, M., Booker, T. R., Czech, L., Gillespie, L., Hateley, S., Kyriazis, C. C., Lang, P. L. M., Leventhal, L., Nogues-Bravo, D., Pagowski, V., Ruffley, M., Spence, J. P., Toro Arana, S. E., Weiß, C. L., & Zess, E. (2022). Genetic diversity loss in the Anthropocene. Science, 377(6613), 1431–1435. 10.1126/science.abn5642

Fabbro, C. D., Scalabrin, S., Morgante, M., & Giorgi, F. M. (2013). An Extensive Evaluation of Read Trimming Effects on Illumina NGS Data Analysis. PLoS ONE, 8(12), e85024. 10.1371/journal.pone.0085024

Frichot, E., Mathieu, F., Trouillon, T., Bouchard, G., & François, O. (2014). Fast and efficient estimation of individual ancestry coefficients. Genetics, 196(4), 973–983. 10.1534/genetics.113.160572

Gaudet, M., Pollegioni, P., Ciolfi, M., Mattioni, C., Cherubini, M., & Beritognolo, I. (2024). Identification of a Unique Genomic Region in Sweet Chestnut (Castanea sativa Mill.) That Controls Resistance to Asian Chestnut Gall Wasp Dryocosmus kuriphilus Yasumatsu. Plants, 13(10), 1355. 10.3390/PLANTS13101355/S1

Geraldes, A., DiFazio, S. P., Slavov, G. T., Ranjan, P., Muchero, W., Hannemann, J., Gunter, L. E., Wymore, A. M., Grassa, C. J., Farzaneh, N., Porth, I., McKown, A. D., Skyba, O., Li, E., Fujita, M., Klápště, J., Martin, J., Schackwitz, W., Pennacchio, C., … Tuskan, G. A. (2013). A 34K SNP genotyping array for Populus trichocarpa: Design, application to the study of natural populations and transferability to other Populus species. Molecular Ecology Resources, 13(2), 306–323. 10.1111/1755-0998.12056

Goudet, J. (2005). Hierfstat, a package for r to compute and test hierarchical F-statistics. Molecular Ecology Notes, 5(1), 184–186. 10.1111/j.1471-8286.2004.00828.x

Gramazio, P., Molina, R., Vilanova, S., Prohens, J., Marrero, Á., Caujapé-Castells, J., & Anderson, G. (2020). Fostering Conservation via an Integrated Use of Conventional Approaches and High-Throughput SPET Genotyping: A Case Study Using the Endangered Canarian Endemics Solanum lidii and S. vespertilio (Solanaceae). Frontiers in Plant Science, 11, 757. 10.3389/fpls.2020.00757

Gremme, G., Steinbiss, S., & Kurtz, S. (2013). GenomeTools: A comprehensive software library for efficient processing of structured genome annotations. IEEE/ACM Transactions on Computational Biology and Bioinformatics, 10(3), 645–656. 10.1109/TCBB.2013.68

Guichoux, E., Garnier-Géré, P., Lagache, L., Lang, T., Boury, C., & Petit, R. J. (2013). Outlier loci highlight the direction of introgression in oaks. Molecular Ecology, 22(2), 450–462. 10.1111/MEC.12125

Guilbaud, R., Biselli, C., Buiteveld, J., Cattivelli, L., Copini, P., Dowkiw, A., Esselink, D., Fricano, A., Guerin, V., Jorge, V., Kelly, Laura. J., Kodde, L., Metheringham, C. L., Pinosio, S., Rogier, O., Segura, V., Spanu, I., Buggs, R., Gonzalez-Martinez, S. C., … Faivre-Rampant, P. (2020, January). Development of a new tool (4TREE) for adapted genome selection in European tree species. https://hal.inrae.fr/hal-02928391

Hemstrom, W., & Jones, M. (2023). snpR: User friendly population genomics for SNP data sets with categorical metadata. Molecular Ecology Resources, 23(4), 962–973. 10.1111/1755-0998.13721

Hoban, S., Hvilsom, C., Aissi, A., Aleixo, A., Bélanger, J., Biala, K., Ekblom, R., Fedorca, A., Funk, W. C., Goncalves, A. L., Gonzalez, A., Heuertz, M., Hughes, A., Ishihama, F., Stroil, B. K., Laikre, L., McGowan, P. J. K., Millette, K. L., O’Brien, D., … da Silva, J. M. (2025). How can biodiversity strategy and action plans incorporate genetic diversity and align with global commitments? BioScience, 75(1), 47–60. 10.1093/biosci/biae106

Homolka, A., Schueler, S., Burg, K., Fluch, S., & Kremer, A. (2013). Insights into drought adaptation of two European oak species revealed by nucleotide diversity of candidate genes. Tree Genetics and Genomes, 9(5), 1179–1192. 10.1007/S11295-013-0627-7/FIGURES/6

Jaramillo-Correa, J. P., Bagnoli, F., Grivet, D., Fady, B., Aravanopoulos, F. A., Vendramin, G. G., & González-Martínez, S. C. (2020). Evolutionary rate and genetic load in an emblematic Mediterranean tree following an ancient and prolonged population collapse. Molecular Ecology, 29(24), 4797–4811. 10.1111/mec.15684

Kastally, C., Niskanen, A. K., Perry, A., Kujala, S. T., Avia, K., Cervantes, S., Haapanen, M., Kesälahti, R., Kumpula, T. A., Mattila, T. M., Ojeda, D. I., Tyrmi, J. S., Wachowiak, W., Cavers, S., Kärkkäinen, K., Savolainen, O., & Pyhäjärvi, T. (2022). Taming the massive genome of Scots pine with PiSy50k, a new genotyping array for conifer research. The Plant Journal, 109(5), 1337–1350. 10.1111/TPJ.15628

Kelly, L. J., Plumb, W. J., Carey, D. W., Mason, M. E., Cooper, E. D., Crowther, W., Whittemore, A. T., Rossiter, S. J., Koch, J. L., & Buggs, R. J. A. (2019). Genes for ash tree resistance to an insect pest identified via comparative genomics (p. 772913). bioRxiv. 10.1101/772913

Kesälahti, R., Kumpula, T. A., Cervantes, S., Kujala, S. T., Mattila, T. M., Tyrmi, J. S., Niskanen, A. K., Rastas, P., Savolainen, O., & Pyhäjärvi, T. (2025). Optimising Exome Captures in Species With Large Genomes Using Species-Specific Repetitive DNA Blocker. Molecular Ecology Resources, 25(3), e14053. 10.1111/1755-0998.14053

Korunes, K. L., & Samuk, K. (2021). pixy: Unbiased estimation of nucleotide diversity and divergence in the presence of missing data. Molecular Ecology Resources, 21(4), 1359– 1368. 10.1111/1755-0998.13326

Kremer, A., Delcamp, A., Lesur, I., Wagner, S., Rellstab, C., Guichoux, E., & Leroy, T. (2024). Whole-genome screening for near-diagnostic genetic markers for four western European white oak species identification. Annals of Forest Science, 81(1), 21. 10.1186/s13595-024-01236-9

Kurtz, S., Narechania, A., Stein, J. C., & Ware, D. (2008). A new method to compute K-mer frequencies and its application to annotate large repetitive plant genomes. BMC Genomics, 9, 517. 10.1186/1471-2164-9-517

Laikre, L., Hoban, S., Bruford, M. W., Segelbacher, G., Allendorf, F. W., Gajardo, G., Rodríguez, A. G., Hedrick, P. W., Heuertz, M., Hohenlohe, P. A., Jaffé, R., Johannesson, K., Liggins, L., MacDonald, A. J., OrozcoterWengel, P., Reusch, T. B. H., Rodríguez-Correa, H., Russo, I.-R. M., Ryman, N., & Vernesi, C. (2020). Post-2020 goals overlook genetic diversity. Science, 367(6482), 1083–1085. 10.1126/science.abb2748

Larue, C., Guichoux, E., Laurent, B., Barreneche, T., Robin, C., Massot, M., Delcamp, A., & Petit, R. J. (2021). Development of highly validated SNP markers for genetic analyses of chestnut species. Conservation Genetics Resources, 13(4), 383–388. 10.1007/S12686-021-01220-9/FIGURES/1

Lefèvre, F., Koskela, J., Hubert, J., Kraigher, H., Longauer, R., Olrik, D. C., Schüler, S., Bozzano, M., Alizoti, P., Bakys, R., Baldwin, C., Ballian, D., Black-Samuelsson, S., Bednarova, D., Bordács, S., Collin, E., De Cuyper, B., De Vries, S. M. G., Eysteinsson, T., … Zariă, I. (2013). Dynamic Conservation of Forest Genetic Resources in 33 European Countries. Conservation Biology, 27(2), 373–384. 10.1111/j.1523-1739.2012.01961.x

Li, H., & Durbin, R. (2009). Fast and accurate short read alignment with Burrows-Wheeler transform. Bioinformatics (Oxford, England), 25(14), 1754–1760. 10.1093/bioinformatics/btp324

Manichaikul, A., Mychaleckyj, J. C., Rich, S. S., Daly, K., Sale, M., & Chen, W.-M. (2010). Robust relationship inference in genome-wide association studies. Bioinformatics (Oxford, England), 26(22), 2867–2873. 10.1093/bioinformatics/btq559

Martin, M. (2011, February). Cutadapt removes adapter sequences from high-throughput sequencing reads. In EMBnet.journal (Vol. 17, Issue 1, pp. 10–12). http://journal.embnet.org/index.php/embnetjournal/article/view/200/479

Mataruga, M., Piotti, A., Daničić, V., Cvjetković, B., Fussi, B., Konnert, M., Vendramin, G. G., & Aleksić, J. M. (2020). Towards the dynamic conservation of Serbian spruce (Picea omorika) western populations. Annals of Forest Science, 77(1), Article 1. 10.1007/s13595-019-0892-1

Mattioni, C., Martin, M. A., Chiocchini, F., Cherubini, M., Gaudet, M., Pollegioni, P., Velichkov, I., Jarman, R., Chambers, F. M., Paule, L., Damian, V. L., Crainic, G. C., & Villani, F. (2017). Landscape genetics structure of European sweet chestnut (Castanea sativa Mill): Indications for conservation priorities. Tree Genetics & Genomes, 13(2), 39. 10.1007/s11295-017-1123-2

Mayol, M., Riba, M., González-Martínez, S. C., Bagnoli, F., de Beaulieu, J.-L., Berganzo, E., Burgarella, C., Dubreuil, M., Krajmerová, D., Paule, L., Romšáková, I., Vettori, C., Vincenot, L., & Vendramin, G. G. (2015). Adapting through glacial cycles: Insights from a long-lived tree (Taxus baccata). New Phytologist, 208(3), 973–986. 10.1111/nph.13496

McKenna, A., Hanna, M., Banks, E., Sivachenko, A., Cibulskis, K., Kernytsky, A., Garimella, K., Altshuler, D., Gabriel, S., Daly, M., & DePristo, M. A. (2010). The Genome Analysis Toolkit: A MapReduce framework for analyzing next-generation DNA sequencing data. Genome Research, 20(9), 1297–1303. 10.1101/GR.107524.110

McKinney, G. J., Waples, R. K., Seeb, L. W., & Seeb, J. E. (2017). Paralogs are revealed by proportion of heterozygotes and deviations in read ratios in genotyping-by-sequencing data from natural populations. Molecular Ecology Resources, 17(4), 656–669. 10.1111/1755-0998.12613

Micheletti, D., Troggio, M., Zharkikh, A., Costa, F., Malnoy, M., Velasco, R., & Salvi, S. (2011). Genetic diversity of the genus Malus and implications for linkage mapping with SNPs. Tree Genetics & Genomes, 7(4), 857–868. 10.1007/s11295-011-0380-8

Milesi, P., Berlin, M., Chen, J., Orsucci, M., Li, L., Jansson, G., Karlsson, B., & Lascoux, M. (2019). Assessing the potential for assisted gene flow using past introduction of Norway spruce in southern Sweden: Local adaptation and genetic basis of quantitative traits in trees. Evolutionary Applications, 12(10), 1946–1959. 10.1111/eva.12855

Milesi, P., Kastally, C., Dauphin, B., Cervantes, S., Bagnoli, F., Budde, K. B., Cavers, S., Fady, B., Faivre-Rampant, P., González-Martínez, S. C., Grivet, D., Gugerli, F., Jorge, V., Kupin, I. L., Ojeda, D. I., Olsson, S., Opgenoorth, L., Pinosio, S., Plomion, C., … Pyhäjärvi, T. (2024). Resilience of genetic diversity in forest trees over the Quaternary. Nature Communications 2024 15:1, 15(1), 1–13. 10.1038/s41467-024-52612-y

Mishra, B., Ulaszewski, B., Meger, J., Aury, J. M., Bodénès, C., Lesur-Kupin, I., Pfenninger, M., Silva, C. D., Gupta, D. K., Guichoux, E., Heer, K., Lalanne, C., Labadie, K., Opgenoorth, L., Ploch, S., Provost, G. L., Salse, J., Scotti, I., Wötzel, S., … Thines, M. (2022). A Chromosome-Level Genome Assembly of the European Beech (Fagus sylvatica) Reveals Anomalies for Organelle DNA Integration, Repeat Content and Distribution of SNPs. Frontiers in Genetics, 12, 691058. 10.3389/FGENE.2021.691058/BIBTEX

Mosca, E., Cruz, F., Gómez-Garrido, J., Bianco, L., Rellstab, C., Brodbeck, S., Csilléry, K., Fady, B., Fladung, M., Fussi, B., san Gömöry, D., González-Martínez, S. C., Grivet, D., Gut, M., Kim Hansen, O., Heer, K., Kaya, Z., Krutovsky, K. V, Kersten, B., … Neale, D. B. (2019). A Reference Genome Sequence for the European Silver Fir (Abies alba Mill.): A Community-Generated Genomic Resource. 10.1534/g3.119.400083

Nocchi, G., Brown, N., Coker, T. L. R., Plumb, W. J., Stocks, J. J., Denman, S., & Buggs, R. J. A. (2022). Genomic structure and diversity of oak populations in British parklands. Plants, People, Planet, 4(2), 167–181. 10.1002/PPP3.10229

Nunziata, A., Ruggieri, V., Petriccione, M., & Masi, L. D. (2020). Single Nucleotide Polymorphisms as Practical Molecular Tools to Support European Chestnut Agrobiodiversity Management. International Journal of Molecular Sciences 2020, Vol. 21, Page 4805, 21(13), 4805. 10.3390/IJMS21134805

Nystedt, B., Street, N. R., Wetterbom, A., Zuccolo, A., Lin, Y. C., Scofield, D. G., Vezzi, F., Delhomme, N., Giacomello, S., Alexeyenko, A., Vicedomini, R., Sahlin, K., Sherwood, E., Elfstrand, M., Gramzow, L., Holmberg, K., Hällman, J., Keech, O., Klasson, L., … Jansson, S. (2013). The Norway spruce genome sequence and conifer genome evolution. Nature 2013 497:7451, 497(7451), 579–584. 10.1038/nature12211

Oggioni, S. D., Rossi, L. M. W., Avanzi, C., Marchetti, M., Piotti, A., & Vacchiano, G. (2024). Drought responses of Italian silver fir provenances in a climate change perspective. Dendrochronologia, 85, 126184. 10.1016/j.dendro.2024.126184

Olsson, S., Dauphin, B., Jorge, V., Grivet, D., Farsakoglou, A. M., Climent, J., Alizoti, P., Faivre-Rampant, P., Pinosio, S., Milesi, P., Scalabrin, S., Bagnoli, F., Scotti, I., Vendramin, G. G., Gonzalez-Martinez, S. C., Fady, B., Aravanopoulus, F. A., Bastien, C., & Alia, R. (2023). Diversity and enrichment of breeding material for resilience in European forests. Forest Ecology and Management, 530, 120748. 10.1016/j.foreco.2022.120748

Olsson, S., Grivet, D., Cattonaro, F., Vendramin, V., Giovannelli, G., Scotti-Saintagne, C., Vendramin, G. G., & Fady, B. (2020). Evolutionary relevance of lineages in the European black pine (Pinus nigra) in the transcriptomic era. Tree Genetics and Genomes, 16(2), 1– 10. 10.1007/S11295-020-1424-8/METRICS

Olsson, S., Pinosio, S., González-Martínez, S. C., Abascal, F., Mayol, M., Grivet, D., & Vendramin, G. G. (2018). De novo assembly of English yew (Taxus baccata) transcriptome and its applications for intra- and inter-specific analyses. 97, 337–345. 10.1007/s11103-018-0742-9

Peace, C., Bassil, N., Main, D., Ficklin, S., Rosyara, U. R., Stegmeir, T., Sebolt, A., Gilmore, B., Lawley, C., Mockler, T. C., Bryant, D. W., Wilhelm, L., & Iezzoni, A. (2012). Development and Evaluation of a Genome-Wide 6K SNP Array for Diploid Sweet Cherry and Tetraploid Sour Cherry. PLOS ONE, 7(12), e48305. 10.1371/JOURNAL.PONE.0048305

Pearman, P. B., Broennimann, O., Aavik, T., Albayrak, T., Alves, P. C., Aravanopoulos, F. A., Bertola, L. D., Biedrzycka, A., Buzan, E., Cubric-Curik, V., Djan, M., Fedorca, A., Fuentes-Pardo, A. P., Fussi, B., Godoy, J. A., Gugerli, F., Hoban, S., Holderegger, R., Hvilsom, C., … Bruford, M. (2024). Monitoring of species’ genetic diversity in Europe varies greatly and overlooks potential climate change impacts. Nature Ecology & Evolution, 8(2), 267–281. 10.1038/s41559-023-02260-0

Pinosio, S., Avanzi, C., Bagnoli, F., Spanu, I., Vendramin, G. G., & Piotti, A. (2025). De Novo transcriptome assembly and annotation of the wild service tree Sorbus torminalis (L.) Crantz. BMC Research Notes, 18(1), 139. 10.1186/s13104-025-07163-w

Pinosio, S., González-Martínez, S. C., Bagnoli, F., Cattonaro, F., Grivet, D., Marroni, F., Lorenzo, Z., Pausas, J. G., Verdú, M., & Vendramin, G. G. (2014). First insights into the transcriptome and development of new genomic tools of a widespread circum-Mediterranean tree species, Pinus halepensis Mill. Molecular Ecology Resources, 14(4), 846–856. 10.1111/1755-0998.12232

Pinosio, S., Marroni, F., Zuccolo, A., Vitulo, N., Mariette, S., Sonnante, G., Aravanopoulos, F. A., Ganopoulos, I., Palasciano, M., Vidotto, M., Magris, G., Iezzoni, A., Vendramin, G. G., & Morgante, M. (2020). A draft genome of sweet cherry (Prunus avium L.) reveals genome-wide and local effects of domestication. The Plant Journal, 103(4), 1420–1432. 10.1111/tpj.14809

Plomion, C., Aury, J. M., Amselem, J., Leroy, T., Murat, F., Duplessis, S., Faye, S., Francillonne, N., Labadie, K., Provost, G. L., Lesur, I., Bartholomé, J., Faivre-Rampant, P., Kohler, A., Leplé, J. C., Chantret, N., Chen, J., Diévart, A., Alaeitabar, T., … Salse, J. (2018). Oak genome reveals facets of long lifespan. Nature Plants, 4(7), 440–452. 10.1038/s41477-018-0172-3

Rellstab, C., & Keller, S. R. (2025). Can We Use Genomic Data to Predict Maladaptation to Environmental Change? Molecular Ecology Resources, 25(4), e14059. 10.1111/1755-0998.14059

Rellstab, C., Zoller, S., Walthert, L., Lesur, I., Pluess, A. R., Graf, R., Bodénès, C., Sperisen, C., Kremer, A., & Gugerli, F. (2016). Signatures of local adaptation in candidate genes of oaks (Quercus spp.) with respect to present and future climatic conditions. Molecular Ecology, 25(23), 5907–5924. 10.1111/MEC.13889

Reutimann, O., Gugerli, F., & Rellstab, C. (2020). A species-discriminatory single-nucleotide polymorphism set reveals maintenance of species integrity in hybridizing European white oaks (Quercus spp.) despite high levels of admixture. Annals of Botany, 125(4), 663–676. 10.1093/AOB/MCAA001

Rey, M. D., Labella-Ortega, M., Guerrero-Sánchez, V. M., Carleial, R., Castillejo, M.Á., Ruggieri, V., & Jorrín-Novo, J. V. (2023). A first draft genome of holm oak (Quercus ilex subsp. Ballota), the most representative species of the Mediterranean forest and the Spanish agrosylvopastoral ecosystem “dehesa.” Frontiers in Molecular Biosciences, 10, 1242943. 10.3389/FMOLB.2023.1242943/FULL

Saleh, D., Chen, J., Leplé, J. C., Leroy, T., Truffaut, L., Dencausse, B., Lalanne, C., Labadie, K., Lesur, I., Bert, D., Lagane, F., Morneau, F., Aury, J. M., Plomion, C., Lascoux, M., & Kremer, A. (2022). Genome-wide evolutionary response of European oaks during the Anthropocene. Evolution Letters, 6(1), 4–20. 10.1002/EVL3.269

Salojärvi, J., Smolander, O. P., Nieminen, K., Rajaraman, S., Safronov, O., Safdari, P., Lamminmäki, A., Immanen, J., Lan, T., Tanskanen, J., Rastas, P., Amiryousefi, A., Jayaprakash, B., Kammonen, J. I., Hagqvist, R., Eswaran, G., Ahonen, V. H., Serra, J. A., Asiegbu, F. O., … Kangasjärvi, J. (2017). Genome sequencing and population genomic analyses provide insights into the adaptive landscape of silver birch. Nature Genetics 2017 49:6, 49(6), 904–912. 10.1038/ng.3862

Scaglione, D., Pinosio, S., Marroni, F., Centa, E. D., Fornasiero, A., Magris, G., Scalabrin, S., Cattonaro, F., Taylor, G., & Morgante, M. (2019). Single primer enrichment technology as a tool for massive genotyping: A benchmark on black poplar and maize. Annals of Botany, 124(4), 543–551. 10.1093/AOB/MCZ054

Shaw, R. E., Farquharson, K. A., Bruford, M. W., Coates, D. J., Elliott, C. P., Mergeay, J., Ottewell, K. M., Segelbacher, G., Hoban, S., Hvilsom, C., Pérez-Espona, S., Ruņģis, D., Aravanopoulos, F., Bertola, L. D., Cotrim, H., Cox, K., Cubric-Curik, V., Ekblom, R., Godoy, J. A., … Grueber, C. E. (2025). Global meta-analysis shows action is needed to halt genetic diversity loss. Nature, 638(8051), 704–710. 10.1038/s41586-024-08458-x

Sollars, E. S. A., Harper, A. L., Kelly, L. J., Sambles, C. M., Ramirez-Gonzalez, R. H., Swarbreck, D., Kaithakottil, G., Cooper, E. D., Uauy, C., Havlickova, L., Worswick, G., Studholme, D. J., Zohren, J., Salmon, D. L., Clavijo, B. J., Li, Y., He, Z., Fellgett, A., McKinney, L. V., … Buggs, R. J. A. (2017). Genome sequence and genetic diversity of European ash trees. Nature, 541(7636), 212–216. 10.1038/nature20786

Stocks, J. J., Metheringham, C. L., Plumb, W. J., Lee, S. J., Kelly, L. J., Nichols, R. A., & Buggs, R. J. A. (2019). Genomic basis of European ash tree resistance to ash dieback fungus. Nature Ecology & Evolution, 3(12), 1686–1696. 10.1038/s41559-019-1036-6

Theraroz, A., Guadaño-Peyrot, C., Archambeau, J., Pinosio, S., Bagnoli, F., Piotti, A., Avanzi, C., Vendramin, G. G., Alía, R., Grivet, D., Westergren, M., & González-Martínez, S. C. (2024). The genetic consequences of population marginality: A case study in maritime pine. Diversity and Distributions, 30(10), e13910. 10.1111/DDI.13910

Tost, M., Grigoriadou-Zormpa, O., Wilhelmi, S., Beissinger, T., Müller, M., Wildhagen, H., Curtu, A. L., & Gailing, O. (2025). Genome-wide association study in European beech (Fagus sylvatica L.) for drought stress traits (p. 2025.04.08.647712). bioRxiv. 10.1101/2025.04.08.647712

Trudić, B., Draškić, G., Provost, G. L., Stojnić, S. D. S., Pilipović, A., & Ivezić, A. (2021). Expression profiles of 11 candidate genes involved in drought tolerance of pedunculate oak (Quercus robur L.). Possibilities for genetic monitoring of the species. Silvae Genetica, 70(1), 226–234. 10.2478/SG-2021-0020

Vajana, E., Andrello, M., Avanzi, C., Bagnoli, F., Vendramin, G. G., & Piotti, A. (2024). Spatial conservation planning of forest genetic resources in a Mediterranean multi-refugial area. Biological Conservation, 293, 110599. 10.1016/j.biocon.2024.110599

Wang, J., Liu, W., Zhu, D., Hong, P., Zhang, S., Xiao, S., Tan, Y., Chen, X., Xu, L., Zong, X., Zhang, L., Wei, H., Yuan, X., & Liu, Q. (2020). Chromosome-scale genome assembly of sweet cherry (Prunus avium L.) cv. Tieton obtained using long-read and Hi-C sequencing. Horticulture Research, 7, 122. 10.1038/s41438-020-00343-8

Weir, B. S., & Cockerham, C. C. (1984). Estimating F-statistics for the analysis of population structure. Evolution; International Journal of Organic Evolution, 38(6), 1358–1370. 10.1111/j.1558-5646.1984.tb05657.x

Xiong, X., Gou, J., Liao, Q., Li, Y., Zhou, Q., Bi, G., Li, C., Du, R., Wang, X., Sun, T., Guo, L., Liang, H., Lu, P., Wu, Y., Zhang, Z., Ro, D. K., Shang, Y., Huang, S., & Yan, J. (2021). The Taxus genome provides insights into paclitaxel biosynthesis. Nature Plants 2021 7:8, 7(8), 1026– 1036. 10.1038/s41477-021-00963-5

Yeaman, S., Hodgins, K. A., Lotterhos, K. E., Suren, H., Nadeau, S., Degner, J. C., Nurkowski, K. A., Smets, P., Wang, T., Gray, L. K., Liepe, K. J., Hamann, A., Holliday, J. A., Whitlock, M. C., Rieseberg, L. H., & Aitken, S. N. (2016). Convergent local adaptation to climate in distantly related conifers. Science, 353(6306), 1431–1433. 10.1126/SCIENCE.AAF7812/SUPPL_FILE/YEAMAN.SM.PDF

Zhao, D., Zhang, Y., Lu, Y., Fan, L., Zhang, Z., Zheng, J., & Chai, M. (2022). Genome sequence and transcriptome of Sorbus pohuashanensis provide insights into population evolution and leaf sunburn response. Journal of Genetics and Genomics, 49(6), 547–558. 10.1016/J.JGG.2021.12.009

Zhou, Q., Karunarathne, P., Andersson-Li, L., Chen, C., Opgenoorth, L., Heer, K., Piotti, A., Vendramin, G. G., Nakvasina, E., Lascoux, M., & Milesi, P. (2024). Recurrent hybridization and gene flow shaped Norway and Siberian spruce evolutionary history over multiple glacial cycles. Molecular Ecology, 33(17), e17495. 10.1111/mec.17495

