## Supplementary Figures for "The FORGENIUS genomic resources: new genotyping tools and genomic data for 23 forest tree species and their Genetic Conservation Units"

**Table of Contents**

| **Figure S1.** Distribution of *Castanea sativa* probes across the *Castanea mollissima* genome in 250 Kb windows. | **Pag. 2** |
| --- | --- |
| **Figure S2.** Distribution of *Fraxinus excelsior* probes across the *Fraxinus excelsior* genome in 250 Kb windows. | **Pag. 2** |
| **Figure S3.** Distribution of *Fagus sylvatica* probes across the *Fagus sylvatica* genome in 250 Kb windows. | **Pag. 3** |
| **Figure S4.** Distribution of *Malus sylvestris* probes across the *Malus sylvestris* genome in 250 Kb windows. | **Pag. 3** |
| **Figure S5.** Distribution of *Pinus halepensis* probes across the *Pinus tabuliformis* genome in 250 Kb windows. | **Pag. 4** |
| **Figure S6.** Distribution of *Prunus avium* probes across the *Prunus avium* genome in 250 Kb windows. | **Pag. 4** |
| **Figure S7.** Distribution of probes across the *Quercus ilex* genome in 250 Kb windows | **Pag. 5** |
| **Figure S8.** Distribution of *Torminalis glaberrima* probes across the *Sorbus pohuashanensis* genome in 250 Kb windows. | **Pag. 5** |
| **Figure S9.** Distribution of *Taxus baccata* probes across the *Taxus chinensis* genome in 250 Kb windows. | **Pag. 6** |
| **Figure S10. A)** Fraction of concordant genotype calls between replicates. **B)** Average sequencing depth at concordant (left) and discordant (right) genotype calls. | **Pag. 6** |
| **Figure S11.** Comparison of genetic variability estimates obtained by stratifying the data based on the species for which the probes were designed. | **Pag. 7** |

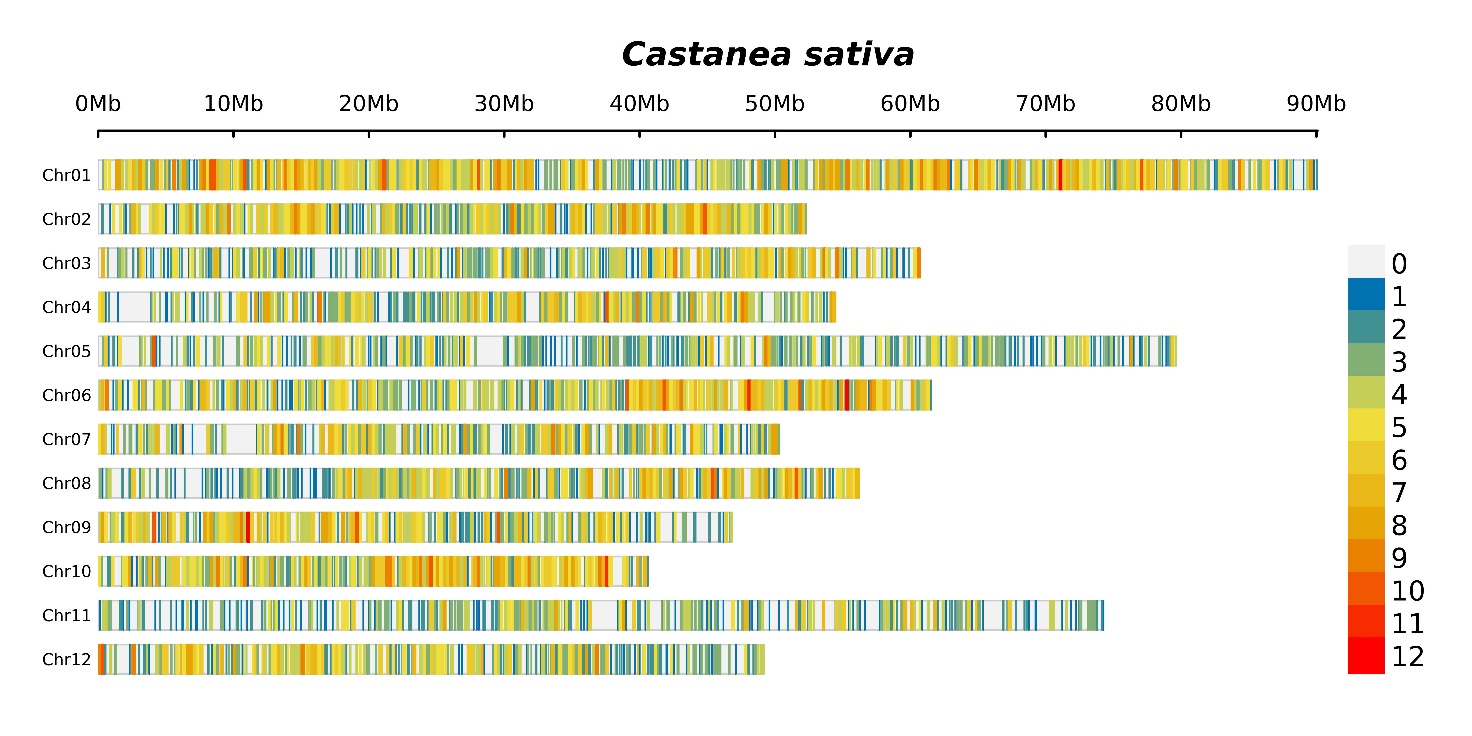

**Figure S1**. Distribution of Castanea sativa probes across the *Castanea mollissima* genome in 250 Kb windows.

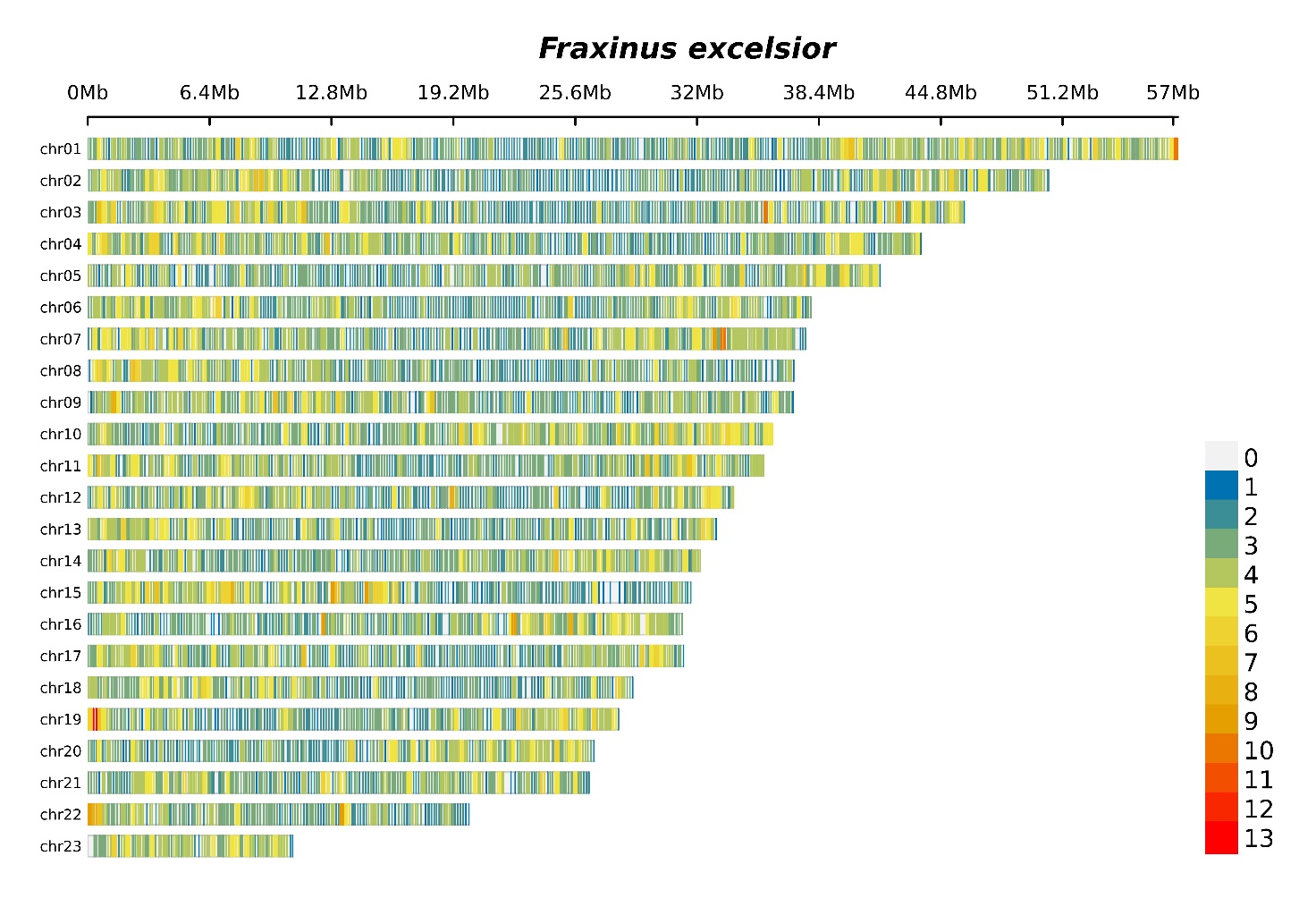

**Figure S2**. Distribution of Fraxinus excelsior probes across the Fraxinus excelsior genome in 250 Kb windows.

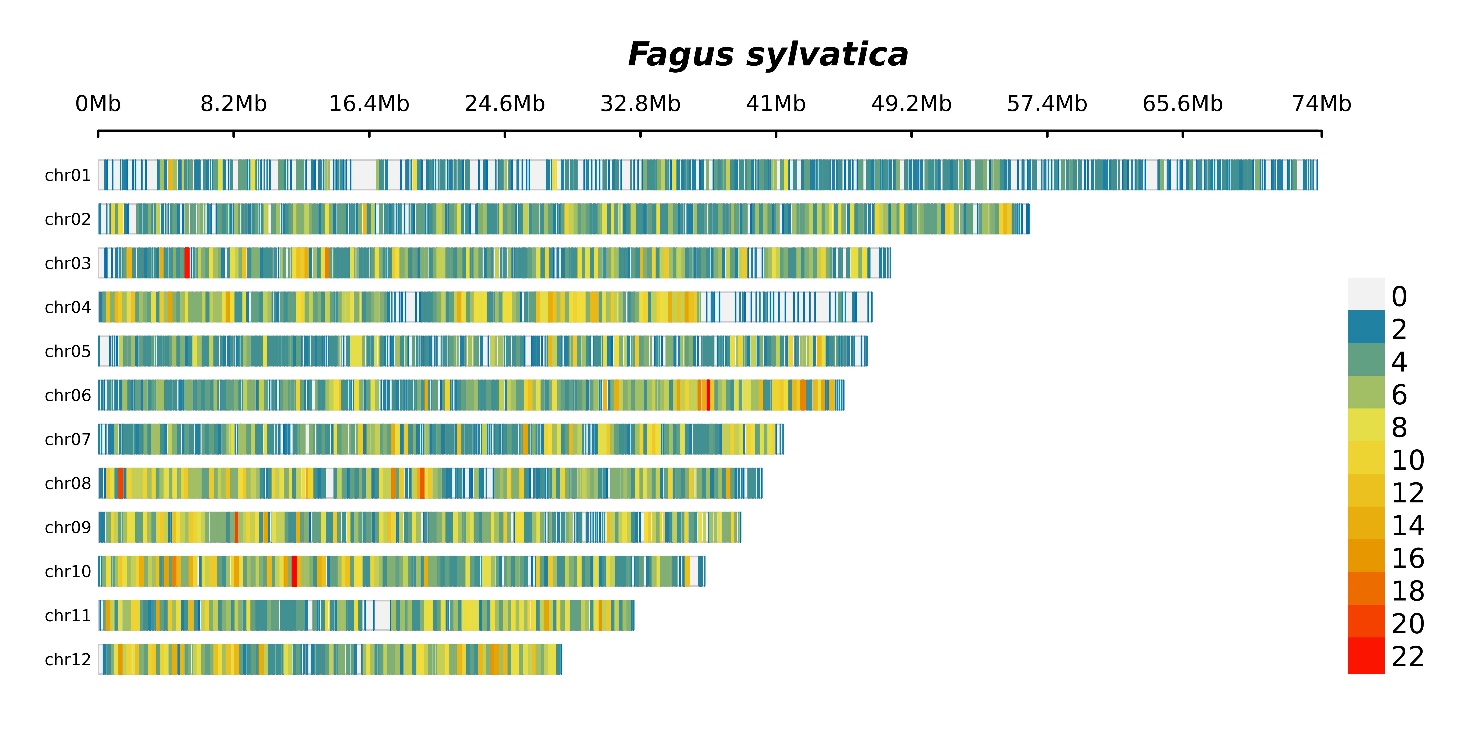
**Figure S3**. Distribution of *Fagus sylvatica* probes across the *Fagus sylvatica* genome in 250 Kb windows.

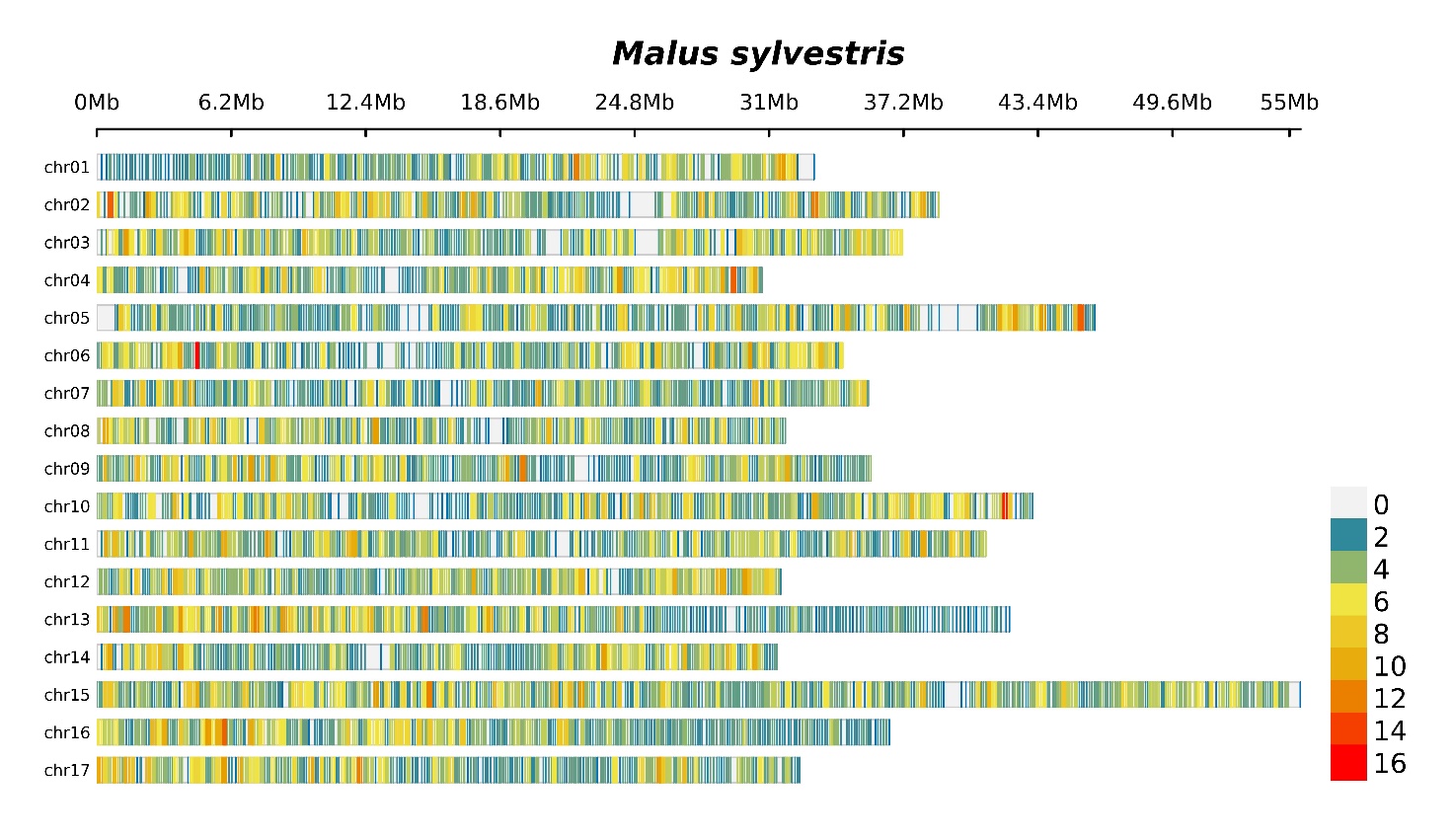
**Figure S4**. Distribution of *Malus sylvestris* probes across the *Malus sylvestris* genome in 250 Kb windows.

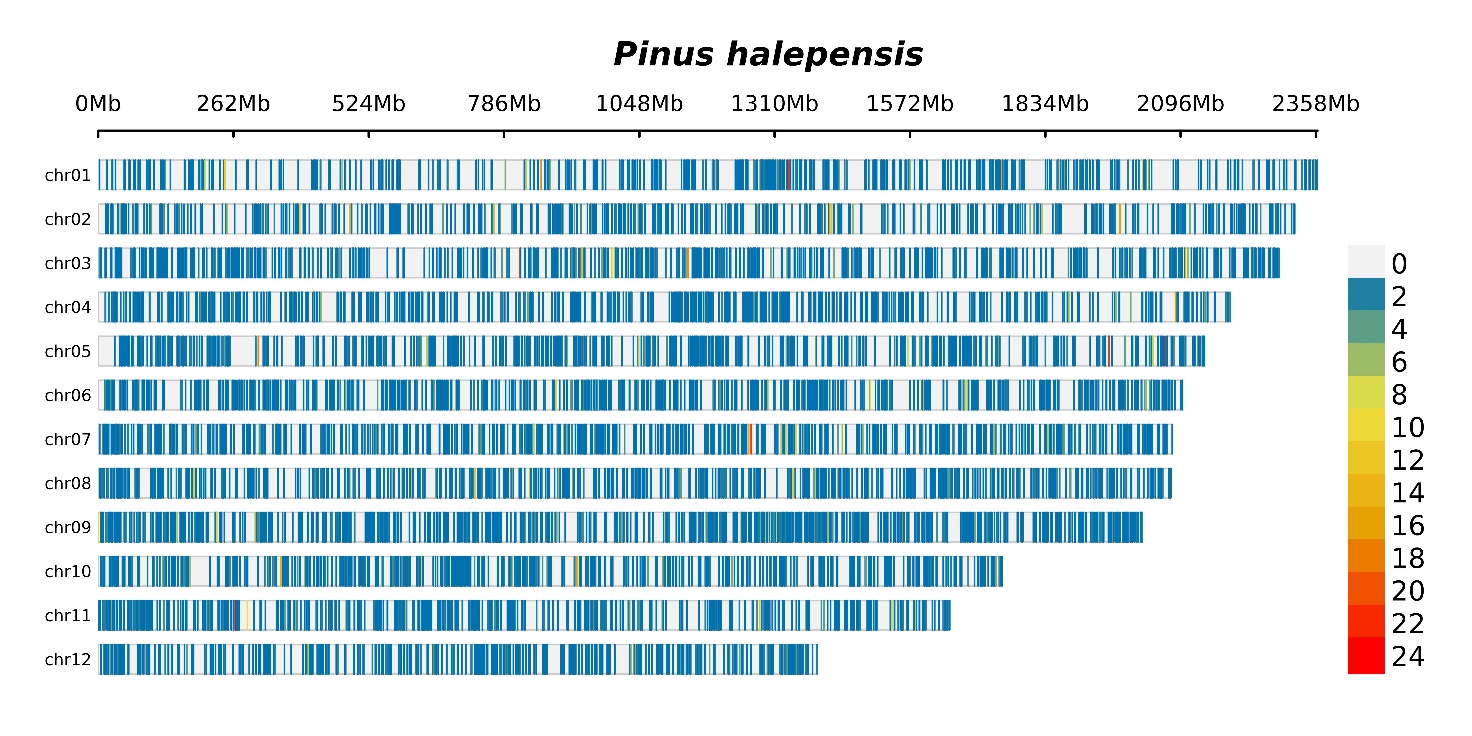
**Figure S5**. Distribution of *Pinus halepensis* probes across the *Pinus tabuliformis* genome in 250 Kb windows.

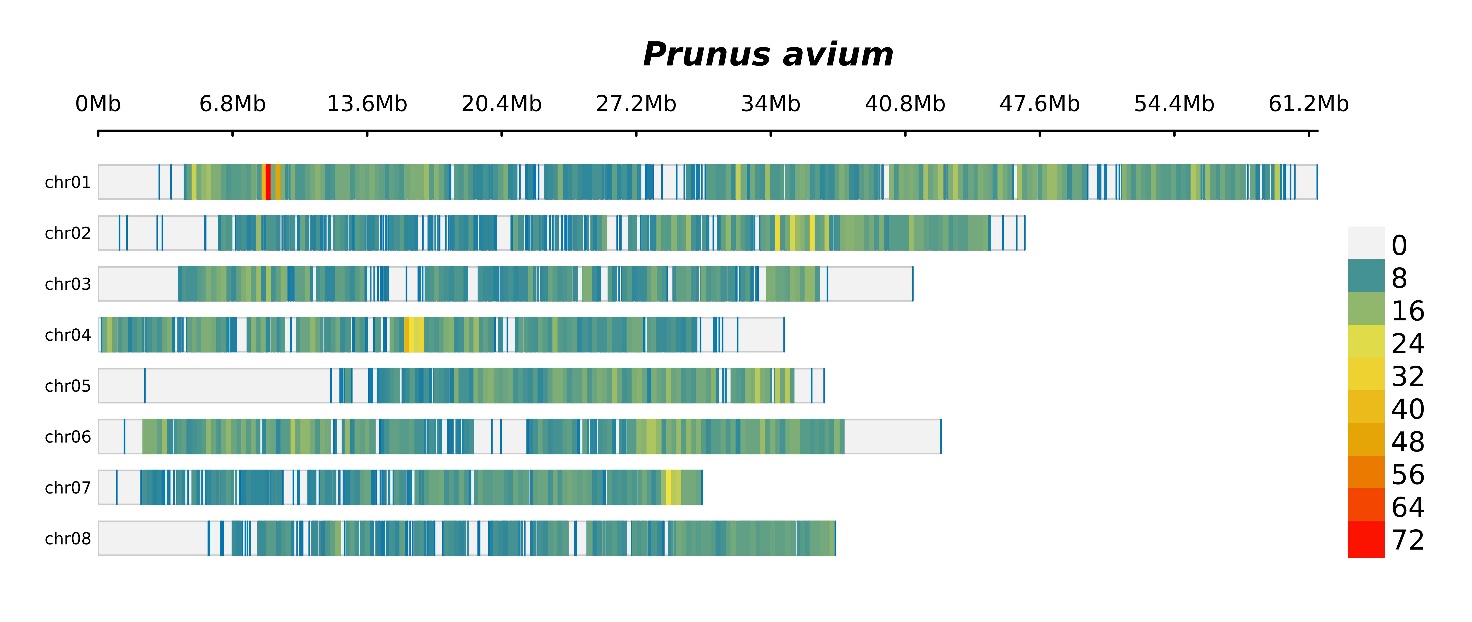
**Figure S6**. Distribution of *Prunus avium* probes across the *Prunus avium* genome in 250 Kb windows.

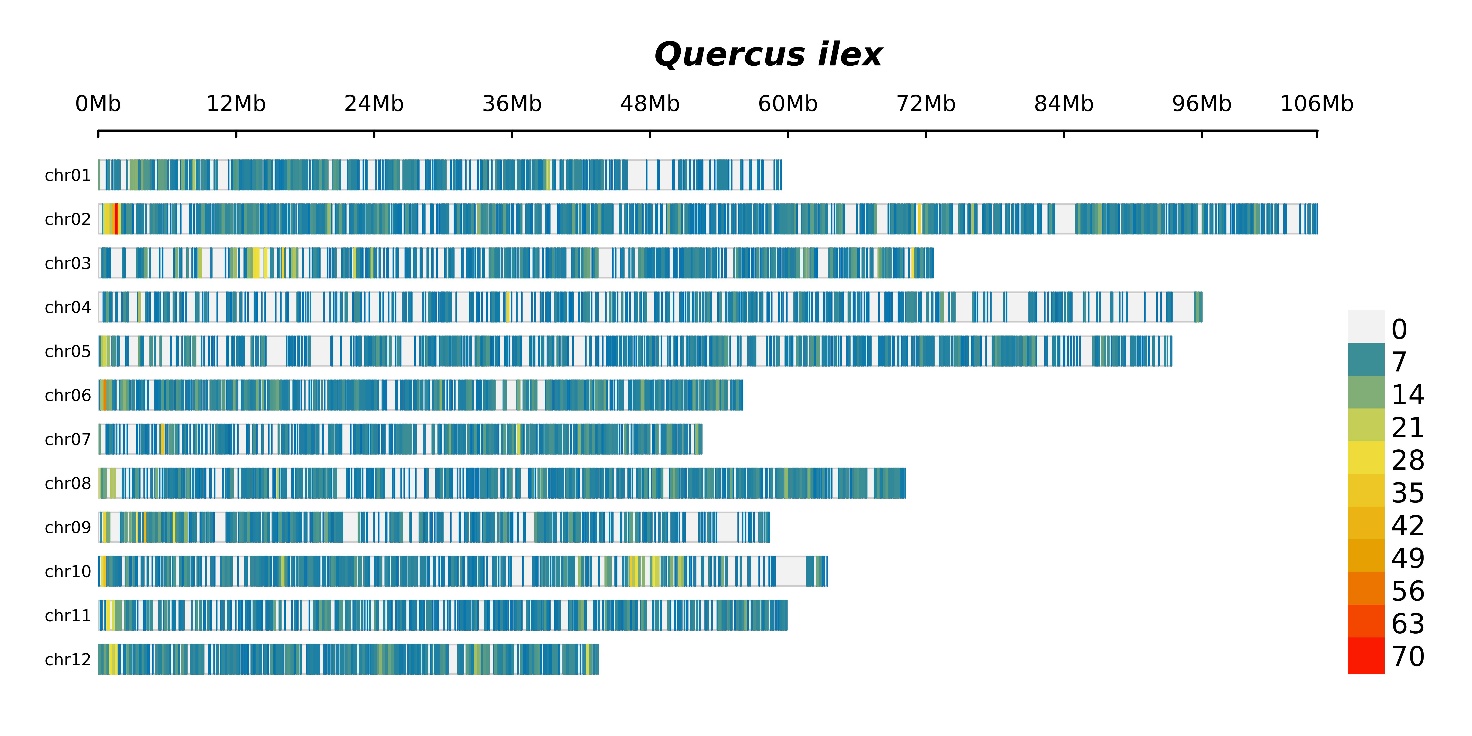
**Figure S7**. Distribution of probes across the *Quercus ilex* genome in 250 Kb windows.

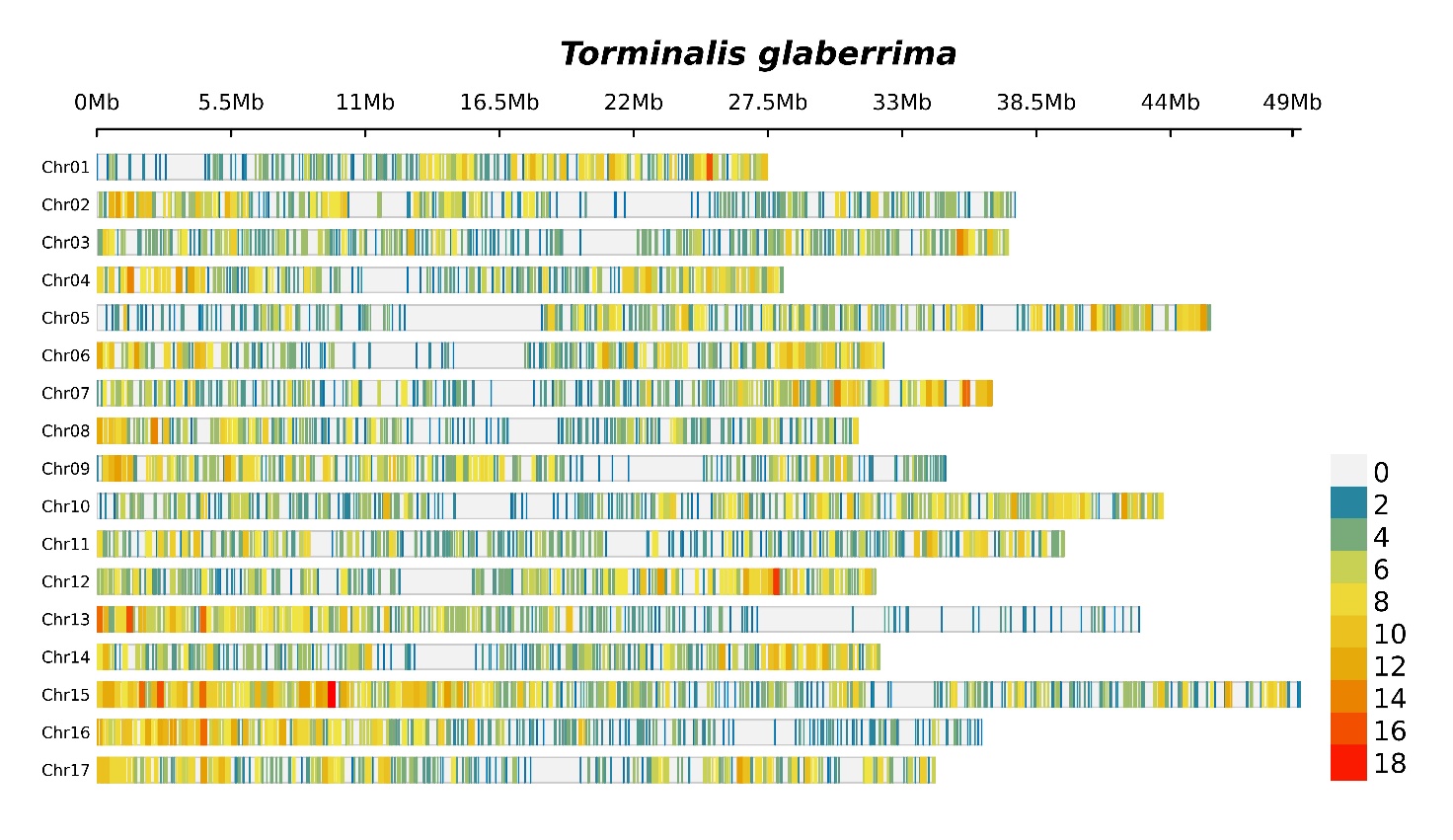
**Figure S8**. Distribution of *Torminalis glaberrima* probes across the *Sorbus pohuashanensis* genome in 250 Kb windows.

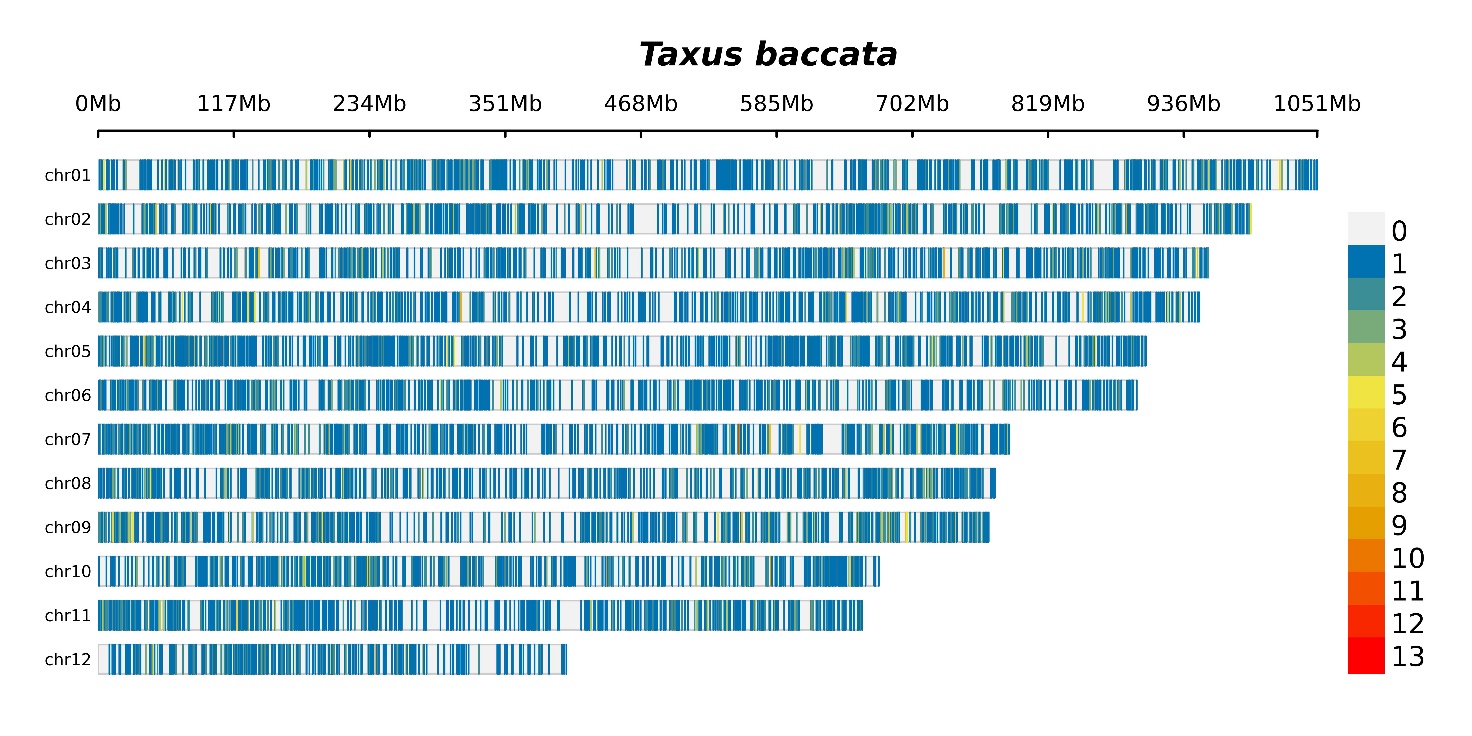
**Figure S9**. Distribution of *Taxus baccata* probes across the *Taxus chinensis* genome in 250 Kb windows.

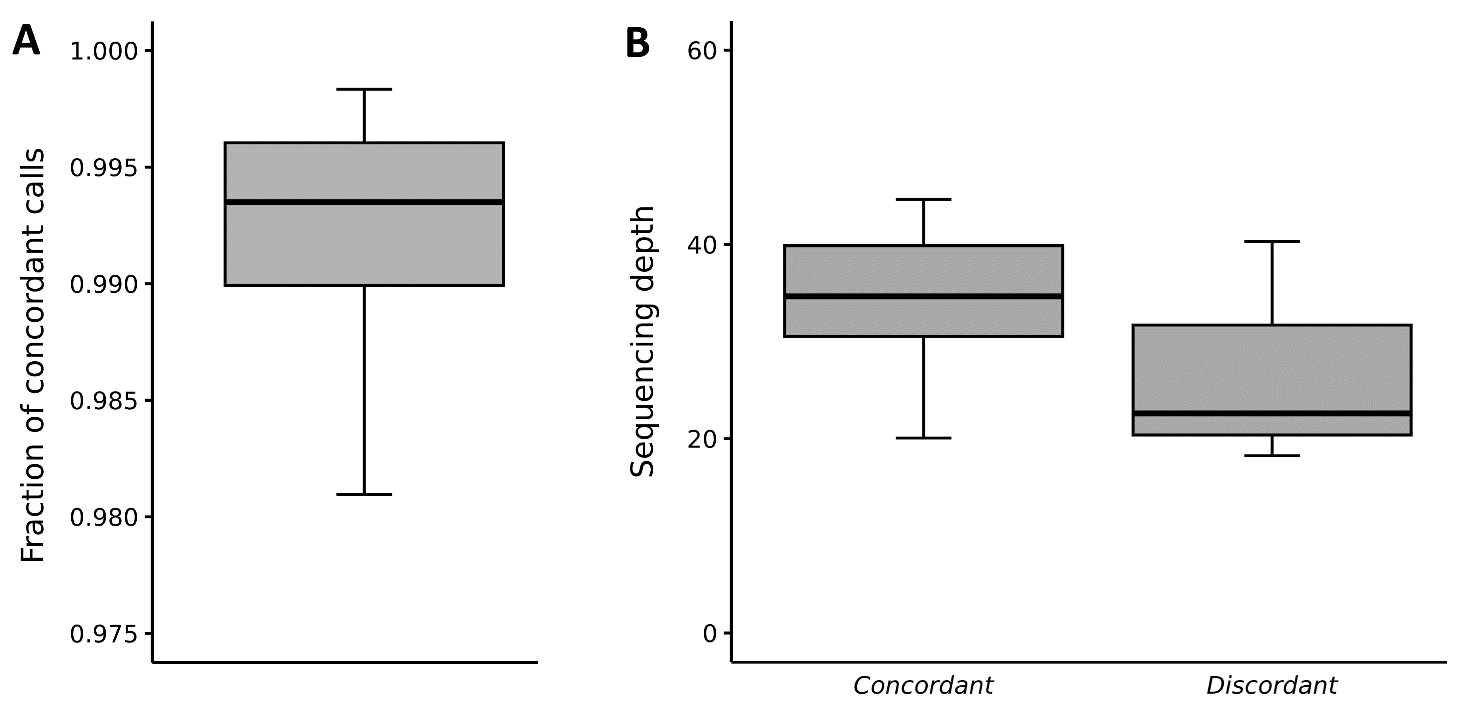

**Figure S10**. **A)** Fraction of concordant genotype calls between replicates. **B)** Average sequencing depth at concordant (left) and discordant (right) genotype calls.

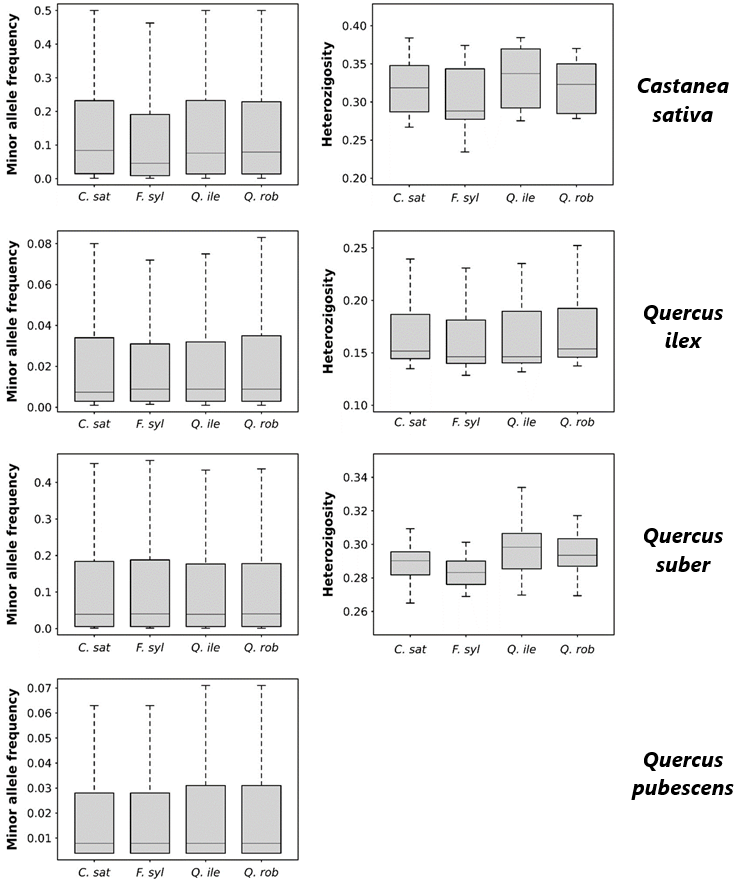

**Figure S11**. Comparison of genetic variability estimates obtained by stratifying the data based on the species for which the probes were designed.
